# Collective interactions augment influenza A virus replication in a host-dependent manner

**DOI:** 10.1101/736108

**Authors:** Kara L. Phipps, Ketaki Ganti, Nathan T. Jacobs, Chung-Young Lee, Silvia Carnaccini, Maria C. White, Miglena Manandhar, Brett E. Pickett, Gene S. Tan, Lucas M. Ferreri, Daniel R. Perez, Anice C. Lowen

## Abstract

Infection with a single influenza A virus (IAV) is only rarely sufficient to initiate productive infection. Here, we exploit both single-cell approaches and whole-animal systems to show that IAV reliance on multiple infection can form an important species barrier to infection. Namely, we find that H9N2 subtype viruses representative of those circulating widely at the poultry-human interface exhibit acute dependence on collective interactions in mammalian systems. This need for multiple infection is greatly reduced in the natural host. Quantification of incomplete viral genomes showed that their complementation accounts for the more moderate reliance on coinfection seen in avian cells, but not the added reliance seen in mammalian cells. This finding suggests an additional form of virus-virus interaction is needed to support infection in mammalian cells. Genetic mapping implicated the PA gene segment as a major driver of this phenotype and quantification of viral RNA synthesis indicated that both replication and transcription were affected. These findings indicate that multiple distinct mechanisms underlie IAV reliance on multiple infection and underscore the importance of virus-virus interactions in IAV infection, evolution and emergence.

## Introduction

Classically, an infectious unit has been defined as a single virus particle which delivers its genome to a cell, initiates the viral reproductive program, and yields progeny viruses. The importance to infection of collective interactions among viruses is being increasingly recognized, however^1–3^. The delivery of multiple viral genomes to a cell allows antagonistic and beneficial interactions to occur, and these interactions have the potential to shape transmission, pathogenicity, and viral evolution.

Recent work has revealed several distinct mechanisms by which multiple viral genomes are co-delivered to a target cell^4–8^. Various mechanisms of direct cell-to-cell spread yield a similar outcome^9–11^. The implications of multiple infection in these diverse systems are still being explored. In a number of cases, however, collective delivery was demonstrated to increase the efficiency of infection relative to free virus particles^4, 5^, or to increase the rate of genetic exchange through recombination^6^.

When distinct variants coinfect, beneficial effects such as compensation of deleterious mutations can increase overall fitness^12–14^. In the case of IAV, several lines of evidence point to a major role in infection for multiplicity reactivation, the process by which segmented genomes lacking one or more functional segments complement each other^15–19^. Conversely, negative interactions can also arise in which deleteriously mutated genes act in a dominant negative fashion. Defective interfering particles, which often potently interfere with the production of infectious progeny from a coinfected cell, are the most extreme example of such antagonism^20–22^. Importantly, multiple infection with identical viral genomes can also alter infection outcomes. Such cooperation was documented for VSV and HIV, where rates of transcription and replication were enhanced with increasing multiplicity of infection (MOI)^23, 24^. Similarly, faster kinetics of virus production were seen at high MOI for poliovirus and IAV^19, 25^. In these instances, it is thought that increased copy number of infecting viral genomes provides a kinetic benefit important in the race to establish infection before innate antiviral responses take hold. Indeed, it has been suggested that multiple infection may be particularly relevant for facilitating viral growth under adverse conditions, such as antiviral drug treatment^3, 26^.

For IAV, an important adverse condition to consider is that of a novel host environment. IAVs occupy a broad host range, including multiple species of wild waterfowl, poultry, swine, humans and other mammals^27, 28^. Host barriers to infection typically confine a given lineage to circulation in one species or a small number of related species^29, 30^. Spillovers occur occasionally, however, and can seed novel lineages. When a novel IAV lineage is established in humans, the result is a pandemic of major public health consequence^31, 32^. The likelihood of successful cross-species transfer of IAV is determined largely by the presence, absence, and compatibility of host factors on which the virus relies to complete its life cycle, and on the viruses’ ability to overcome antiviral defenses in the novel host^33–35^.

Our objective herein was to assess the degree to which IAV relies on the delivery of multiple viral genomes to a cell to ensure production of progeny. In particular, we sought to determine whether this phenotype varies with host species. We therefore examined the multiplicity dependence of one human and a panel of avian-origin viruses in multiple host systems. Results from all virus/cell combinations tested confirm prior reports that cells multiply-infected with IAV produce more viral progeny than singly-infected cells. Importantly, however, the extent to which viral progeny production is concentrated within the multiply-infected fraction of a cell population varies greatly with virus-host context. Two poultry-adapted H9N2 viruses (A/guinea fowl/HK/WF10/99 (GFHK99) and A/quail/HK/A28945/88 (QaHK88)) exhibit an acute dependence on multiple infection in mammalian systems that is greatly diminished in natural host systems. This strong dependence on multiple infection is not seen for the human strain, influenza A/Netherlands/602/2009 (H1N1) (NL09) virus, or for an IAV from wild birds, influenza A/mallard/199106/99 (H3N8) (MaMN99) virus. The host-specific dependence of GFHK99 virus on multiple infection is driven by its PA gene segment. In line with this finding, both bulk and single-cell measurements of viral RNA showed that polymerase activity in mammalian cells is enhanced with multiple infection. A need for complementation of incomplete viral genomes partially accounts for this cooperative effect. Importantly, however, single-cell data indicate that additional multiplicity-dependent mechanisms support RNA synthesis by the GFHK99 virus in mammalian cells. Thus, our data point to an important role for multiple infection in determining the potential for IAV replication in diverse hosts.

## Results

### Degree of multiplicity dependence varies with strain and cell type

To evaluate the extent to which IAV relies on multiple infection for productive infection, we initially used reassortment and coinfection as readouts. While singly infected cells yield only parental progeny, reassortment is highly efficient within coinfected cells^36^. Monitoring reassortment therefore indicates the proportion of progeny in a bulk virus population that were produced from multiply infected cells. To ensure accurate quantification of reassortment, coinfections were performed under single-cycle conditions with homologous viruses that differ only by a silent mutation in each segment and the presence of either an HA or HIS epitope tag fused to the HA protein. The silent mutations allow identification of reassortant viruses. The epitope tags enable quantification of cellular coinfection across a range of MOIs. Homologous coinfection partners, termed WT and VAR, were generated in NL09, MaMN99 and GFHK99 strain backgrounds.

Coinfection and reassortment were examined in canine MDCK and chicken DF-1 cells (**Figure 1**). In MDCK cells, GFHK99 viruses showed a near linear relationship between total HA^+^ cells and dual-HA^+^ cells, suggesting a strict dependence on multiple infection for HA expression (**Figure 1A**). By contrast, HA expression resulting from infection with a single strain was more common for GFHK99 in DF-1 cells, MaMN99 in MDCK cells and NL09 in MDCK cells (**Figure 1A**). To give a quantitative measure of the differences among these virus/cell pairings, we determined the degree of linearity from the regression models of each dataset (**Figure 1B**). The more dependent expression of *any* HA protein is on coinfection, the more linear the relationship between dual-HA^+^ cells and HA^+^ cells becomes. Conversely, a relationship closer to the Poisson expectation indicates less dependence on coinfection. Among the viruses tested, we found that only GFHK99 virus in MDCK cells exhibits strong linearity in the relationship between HA^+^ and dual-HA^+^ cells (**Figure 1B**).

**Figure 1.**
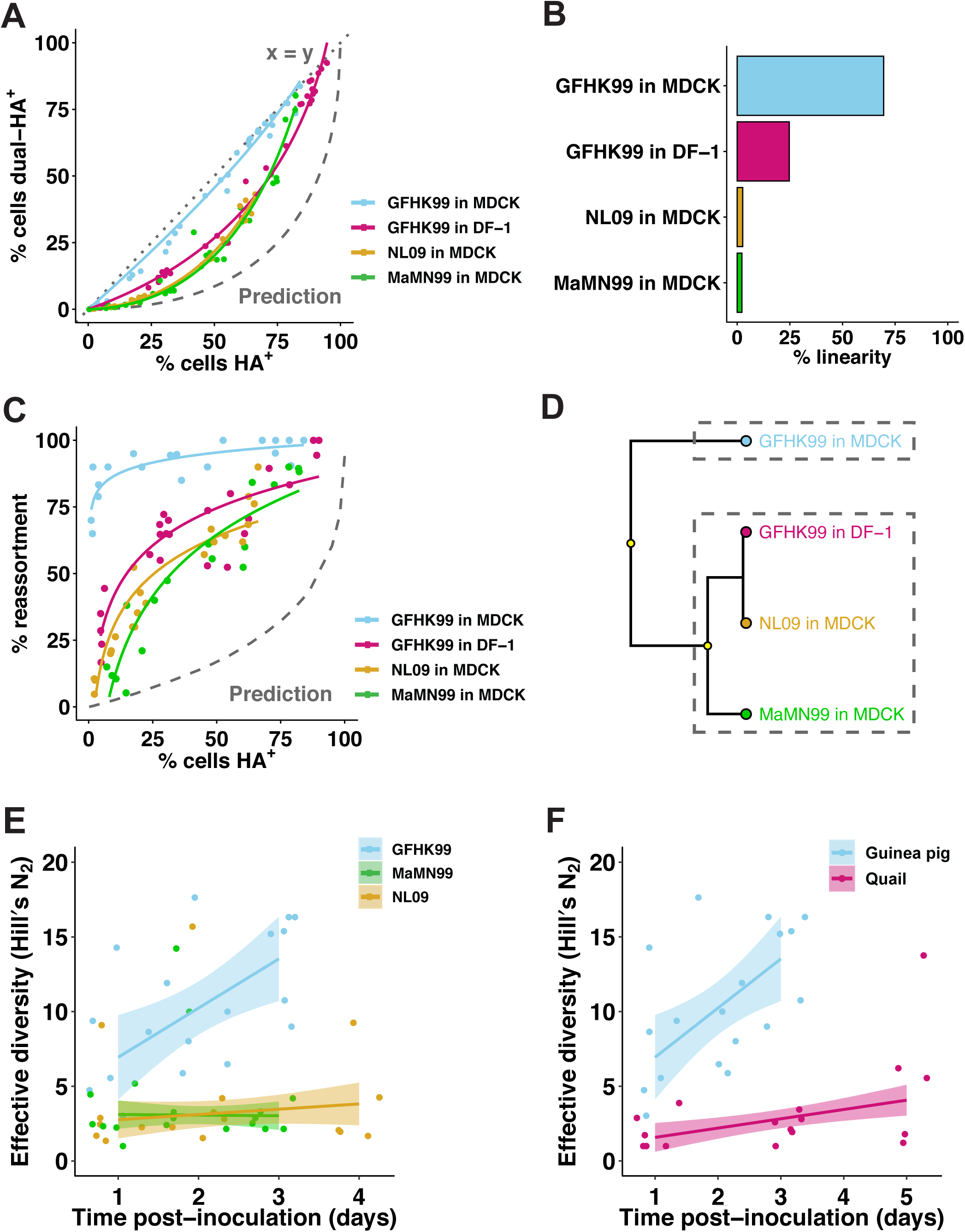
Coinfection and reassortment frequencies indicate that IAV multiplicity dependence varies with virus strain and host species. A–D) MDCK or DF-1 cells were coinfected with homologous WT and VAR viruses of either GFHK99, MaMN99, or NL09 strain backgrounds at a range of MOIs. The relationship between % cells HA^+^ and % cells dual-HA^+^ (A) varies with strain and cell type, resulting in curves of differing % linearity (B). GFHK99, MaMN99, and NL09 viruses exhibit different reassortment levels in MDCK cells, but all show high reassortment relative to a theoretical prediction in which singly infected and multiply infected cells have equivalent burst sizes (C). GFHK99 virus reassortment levels differ in MDCK and DF-1 cells, but again reassortment in DF-1 cells remains high relative to the theoretical prediction in which multiple infection confers no advantage (C). Clustering analysis of reassortment and HA co-expression regression models determines that GFHK99 virus exhibits unique behavior in MDCK cells compared to DF-1 cells or other viruses in MDCK cells (D). Yellow circles indicate nodes with >95% bootstrap support. In guinea pigs (n=6), GFHK99 WT and VAR_1_ viruses exhibit higher reassortment than MaMN99 or NL09 WT and VAR viruses, as indicated by increased genotypic diversity (E). The GFHK99 WT and VAR_1_ viruses exhibit higher reassortment in guinea pigs than in quail (n=5) (F). Guinea pig data shown in panels E and F are the same. NL09 virus reassortment data shown in (C) were reported previously^76^. Shading represents 95% CI.

In line with observed levels of coinfection, GFHK99 virus exhibits high levels of reassortment in MDCK cells, indicating that nearly all progeny virus is produced from WT-VAR_1_ coinfected cells (**Figure 1C**). MaMN99 or NL09 viruses infecting MDCK cells show relatively lower levels of reassortment. Moreover, reassortment of GFHK99 viruses is markedly reduced in DF-1 cells compared to MDCK cells (**Figure 1C**). These results clearly reveal differing degrees of multiplicity dependence for different virus/cell pairings and therefore indicate that multiple infection dependence is determined through virus-host interactions.

That all virus-cell pairings tested show evidence of multiplicity dependence is highlighted by comparison of the experimental reassortment data to a theoretical prediction (**Figure 1C**). This theoretical prediction was published previously^18^ and is derived from a computational model in which the number of viral progeny produced by an infected cell is constant, irrespective of the number of input viral genomes. Because singly- and multiply-infected cells make equivalent numbers of progeny in this model, reassortment is predicted to increase only gradually at low levels of infection where coinfection is relatively rare. The experimental observation that reassortment reaches high levels much more rapidly than expected indicates that viral progeny production is enriched in the proportion of the infected cell population that is multiply infected.

Frequent reassortment and the linear relationship between overall HA expression and HA co-expression provide two measures of multiplicity dependence for a given virus-cell pairing. To summarize these two measures and analyze the relationships between the virus-cell pairings tested, we conducted a hierarchical clustering analysis using the parameters of the regression models shown in **Figure 1A** and **1C**. MaMN99 and NL09 viruses in MDCK cells, as well as GFHK99 virus in DF-1 cells, represent one cluster, while GFHK99 virus in MDCK cells partitioned into its own cluster (**Figure 1D**). Coinfection dependence, therefore, is evident in all virus-cell pairings, but particularly strong for GFHK99 virus in MDCK cells.

### Strain- and host-specific phenotypes are also evident *in vivo*

To determine whether host-dependent reliance on multiple infection extended to *in vivo* models, we performed coinfections in guinea pigs and quail. GFHK99, MaMN99, and NL09 viruses were tested in guinea pigs and GFHK99 virus was examined in quail. To ensure use of comparable effective doses for each virus/host pairing, the 50% infectious dose (ID_50_) of each virus mixture was first determined experimentally in the animal models used. Guinea pigs were then infected intranasally with 10^2^ GPID_50_ of each WT/VAR mixture and nasal washes were collected daily. Quail were infected with 10^2^ QID_50_ of the GFHK99 virus mixture via an oculo-naso-tracheal route and tracheal swabs were collected daily. To evaluate the frequency of reassortment, plaque isolates from these upper respiratory samples were genotyped for each animal on each day. Because multicycle replication *in vivo* allows the propagation of reassortants, analysis of genotypic diversity rather than percent reassortment is more informative for these experiments. Thus, the effective diversity (Hill’s N_2_) was calculated for each dataset and plotted as a function of time post-inoculation (**Figure 1E, 1F**). The viruses collected from GFHK99-infected guinea pigs show much higher genotypic diversity throughout the course of infection than viruses isolated from MaMN99- or NL09-infected guinea pigs (**Figure 1E**) or GFHK99-infected quail (**Figure 1F**). These data indicate that the virus-host interactions which determine dependence on multiple infection in cell culture extend to *in vivo* infection.

### Multiple infection enhances viral growth

The abundant reassortment observed with GFHK99 viruses in mammalian systems suggests that multiple infection plays a major role in determining the productivity of an infected cell. We therefore hypothesized that increasing MOI would augment the viral output of infected cells and that the magnitude of this effect would be greater for GFHK99 virus in MDCK cells than for GFHK99 virus in DF-1 cells or MaMN99 and NL09 viruses in MDCK cells. To test this prediction, we infected over a range of MOIs and then measured PFU produced per cell under single-cycle conditions. The input MOIs for this series of experiments were based on PFU as determined in MDCK cells. Which MOI conditions were saturating was determined empirically by flow cytometry in MDCK and DF-1 cells (**Supplementary Figure 1**). MOIs < 1 PFU/cell were found to be non-saturating. Under these non-saturating conditions, increasing MOI resulted in accelerated viral growth and higher burst size for all virus/cell pairings (**Figure 2A–D**). As predicted, however, increasing the MOI of GFHK99 virus in MDCK cells resulted in a further enhancement of viral amplification (**Figure 2G**). Thus, the benefit conferred by multiple infection was greater for GFHK99 virus in MDCK cells compared to the other virus-cell pairings tested.

**Figure 2.**
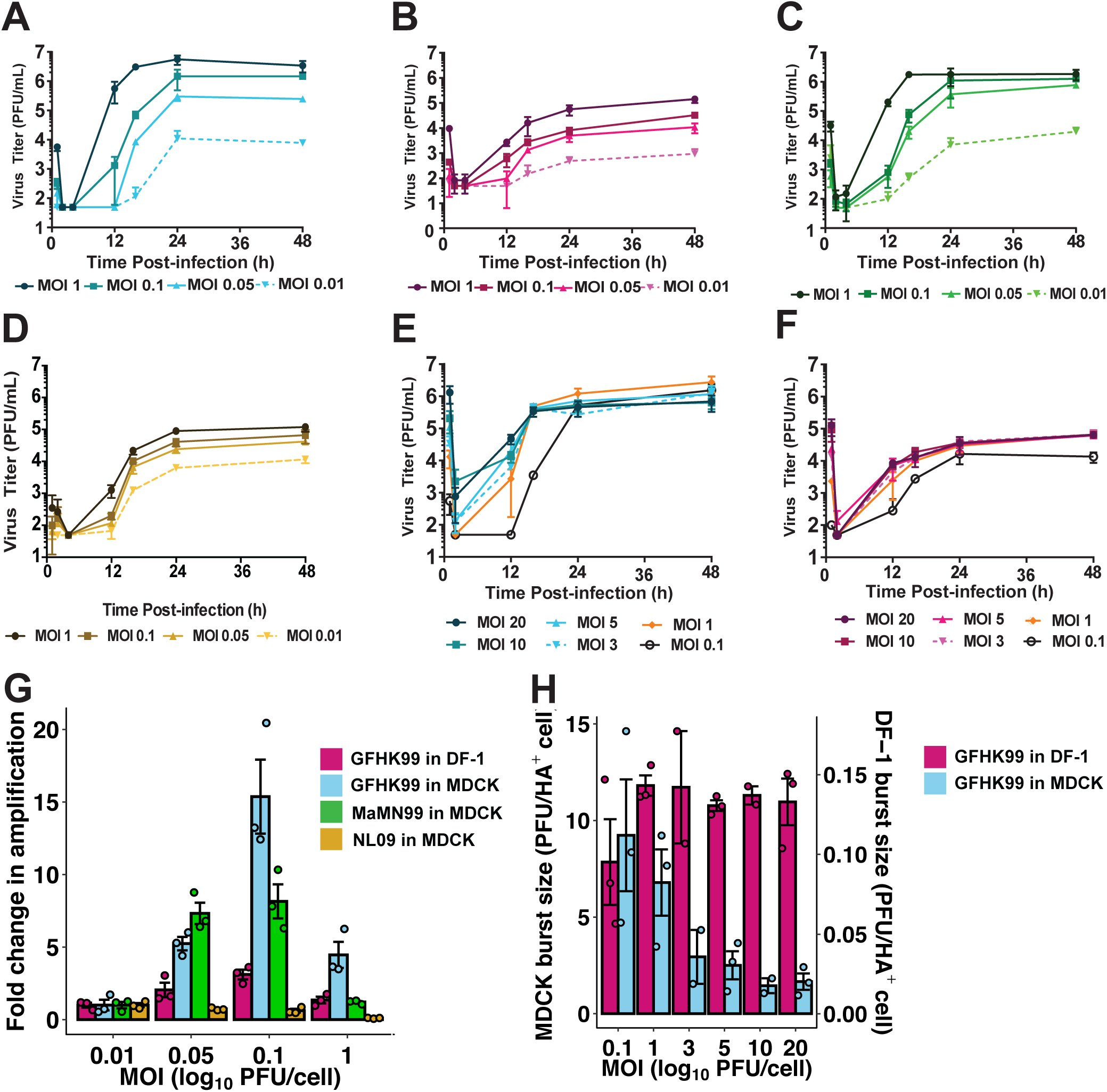
Increasing MOI increases viral productivity at sub-saturating, but not saturating MOIs. MDCK or DF-1 cells were infected under single-cycle conditions at a range of MOIs in triplicate wells for each MOI. A–F) Viral titers observed at the indicated MOIs are plotted against time post-infection for GFHK99 virus in MDCK cells (A), GFHK99 virus in DF-1 cells (B), MaMN99 virus in MDCK cells (C), NL09 virus in MDCK cells (D), GFHK99 virus in MDCK cells (E), and GFHK99 virus in DF-1 cells (F). G) Fold change in amplification (maximum PFU output / PFU input) relative to the MOI=0.01 PFU per cell condition is plotted for each virus-cell pairing. H) Burst size, calculated as maximum PFU output / number of HA^+^ cells detected by flow cytometry, is plotted for each virus-cell pairing tested in the higher MOI range. MOIs shown are in units of PFU per cell, as determined in MDCK cells. Error bars represent mean ± standard error.

We reasoned that the cooperative effect observed might result from i) complementation of incomplete viral genomes or ii) a benefit of increased viral genome copy number per cell. In an effort to differentiate between these possibilities, we measured growth of GFHK99 virus in MDCK and DF-1 cells infected over range of saturating MOIs. Note that plaque assays in MDCK cells underestimate the infectious potential of GFHK99 virus, and increasing viral MOI beyond 1 PFU/cell did not lead to increased levels of HA+ cells, indicating saturation (**Supplementary Figure 1**). Under these conditions, incomplete viral genomes are unlikely to be prevalent and any benefit of increasing MOI would be attributable to increasing genome copy numbers per cell. In both cell types, MOIs between 1 and 20 PFU per cell result in similar peak viral titers (**Figure 2E, F, H**). This saturation of cooperation at higher MOIs suggests diminishing returns from additional genome copies above a certain threshold. Whether this threshold is imposed by a need for complementation or another mechanism sensitive to saturation remained unclear.

### Multiple infection enhances viral RNA replication

To test whether multiple infection was beneficial to the virus at the level of genome replication, we measured WT viral RNA synthesis in the absence and presence of increasing amounts of a homologous coinfecting virus. We also broadened the panel of viruses examined to include two additional strains: A/duck/HK/448/78 (H9N2) (DkHK78), a wild bird isolate that predates the establishment of H9N2 viruses in poultry, and A/quail/HK/88 (H9N2) (Qa/HK/88), an early isolate of the H9N2 poultry lineage^37^. To evaluate host specificity, we measured WT viral RNA synthesis in both mammalian and avian cells (**Figure 3**). Cells were infected with low MOI of WT virus to ensure receipt of a single copy of the WT virus genome. Concurrently, cells were infected with increasing doses of a VAR virus. Droplet digital PCR (ddPCR) was then used to quantify replication of WT genomes using primer/probe sets that do not detect VAR templates.

**Figure 3.**
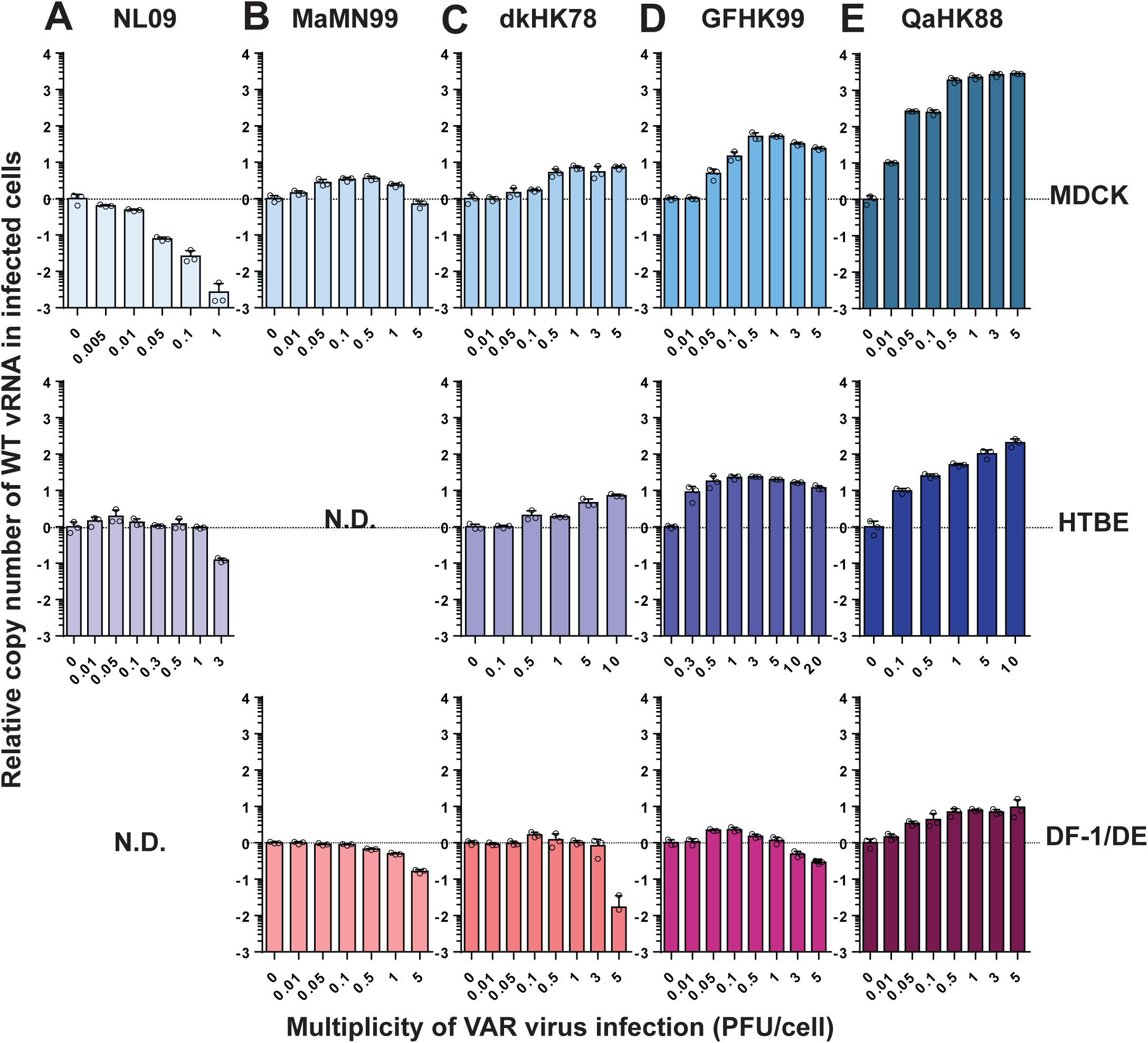
Coinfection enhances GFHK99 vRNA synthesis in a dose and host dependent manner. Cells were coinfected with WT virus and increasing doses of VAR virus. WT virus MOI was 0.05 PFU per cell in NHBE cells and 0.005 PFU per cell in all other cell types. The fold change in WT vRNA copy number, relative to that detected in the VAR virus, is plotted for NL09 virus in MDCK and NHBE cells (A), MaMN99 virus in MDCK and duck embryo (DE) cells (B), dkHK78 virus in MDCK, NHBE, and DF-1 cells (C), GFHK99 virus in MDCK, NHBE, and DF-1 cells (D) and QaHK88 virus in MDCK, NHBE, and DF-1 cells (E). n=3 cell culture dishes per condition. Error bars represent mean ± standard deviation.

The results for NL09 and MaMN99 viruses show little to no increase in WT vRNA production with VAR virus addition, in all cell types tested (**Figure 3A** and B). DkHK78 virus exhibits an intermediate phenotype, with up to 7-fold increases in WT vRNA levels seen with VAR virus addition in MDCK and NHBE cells. For this virus in avian DF-1 cells, only a 1.6-fold increase is observed (**Figure 3C**). GFHK99 and QaHK88 viruses show much stronger enhancement of WT vRNA replication with VAR virus coinfection: in MDCK and NHBE cells, respectively, GFHK99 virus show increases of 60-fold and 20-fold, while QaHK88 virus shows increases of more than 2000-fold and 200-fold. In contrast, these two poultry isolates show markedly less enhancement in WT vRNA replication with VAR coinfection in avian DF-1 cells (**Figure 3D** and E). Of note, addition of VAR at high doses suppresses WT vRNA production in several virus/cell pairings, likely indicating competition for limited resources in the cell. This antagonistic interaction is stronger in systems where little benefit of coinfection is detected.

In summary, for both GFHK99 and QaHK88 viruses, the introduction of coinfecting VAR virus reveals a cooperative effect acting at the level of RNA synthesis, which is particularly potent in mammalian cells. That the replication of NL09 and MaMN99 viruses is not detectably enhanced by the addition of VAR virus suggests a reduced reliance on multiple infection, consistent with reassortment and dual-HA positivity data presented above.

### The viral PA segment is a major determinant of multiple infection dependence

To identify viral genetic determinants of multiple infection dependence, we mapped segments responsible for the high reassortment phenotype of GFHK99 virus in MDCK cells. One or more genes from GFHK99 virus was placed into a MaMN99 virus background. For each chimeric genotype, WT and VAR strains were generated to allow tracking of homologous reassortment. Coinfections were performed in MDCK cells and HA expression and reassortment were measured as in **Figure 1**. Two viruses are similar to the parental GFHK99 virus in the levels of dual HA positivity observed: MaMN99:GFHK99 3PNP and MaMN99:GFHK99 PA (**Figure 4A– B**). Quantification of reassortment revealed that several chimeric viruses reassort at a higher frequency than MaMN99 parental strains, but that the MaMN99:GFHK99 3PNP and MaMN99:GFHK99 PA viruses show the highest reassortment, comparable to that seen for the parental GFHK99 genotype (**Figure 4C-D**). A clustering analysis similar to the one shown in **Figure 1D** revealed that only these two genotypes cluster with GFHK99 parental virus, while each of the other genotypes clusters with MaMN99 virus (**Figure 4E**). Thus, while other viral genes may contribute, PA is the primary genetic determinant of the high reassortment exhibited by GFHK99 virus in MDCK cells and defines a need for cooperation between coinfecting viruses. To verify the contribution of the PA segment by a second approach, we evaluated the impact of VAR virus coinfection on WT vRNA synthesis for the MaMN99:GFHK99 PA chimeric strain in MDCK and DF-1 cells. The results confirm that the PA segment of GFHK99 virus increases reliance on multiple infection, specifically in mammalian cells (**Supplementary Figure 2**).

**Figure 4.**
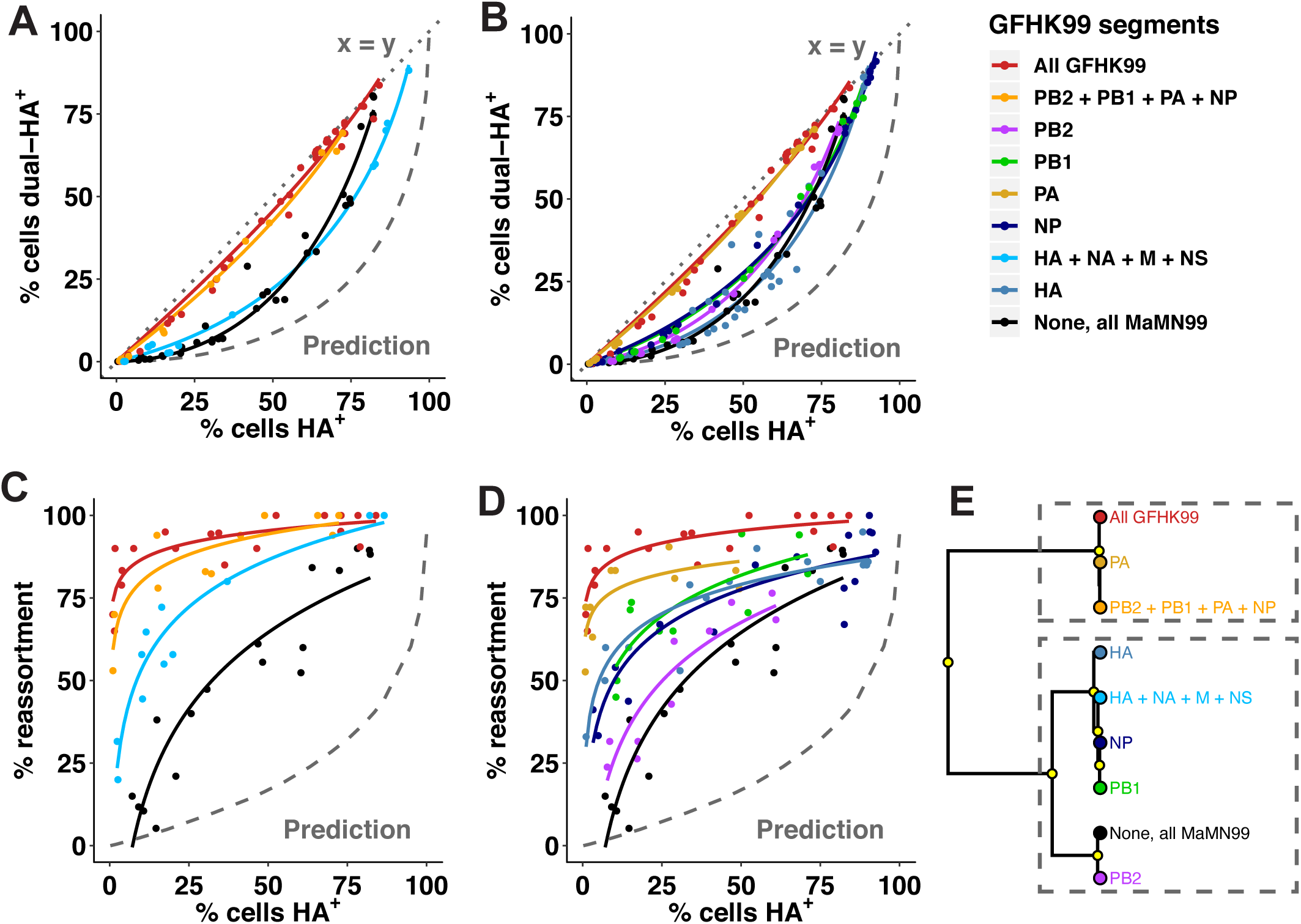
Coinfection and reassortment of chimeric viruses reveal a major role for the viral PA gene segment. Reverse genetics was used to place one or more genes from GFHK99 virus into a MaMN99 background. Coinfections with homologous WT and VAR strains were performed in MDCK cells as in Figure 1. The relationship between % cells HA positive and % cells dually HA positive (A, B) and reassortment levels (C, D) are shown for each genotype. Experimental results are compared to a theoretical prediction based on Poisson statistics and in which singly infected and multiply infected cells have equivalent burst sizes (Prediction). Clustering analysis of the regression models shown in (A-D) is used to partition the behavior of each chimeric genotype into one of two clusters, denoting closer similarity to GFHK99 or MaMN99 (E). Yellow circles indicate nodes with >95% bootstrap support. Data shown for GFHK99 and MaMN99 viruses are the same as those displayed in Figure 1.

### Multiple infection accelerates viral replication and transcription

Because mapping of the GFHK99 high reassortment phenotype implicated the viral PA, we tested the effects of multiple infection on viral replication and transcription by measuring vRNA and mRNA over time following low or high MOI infection (**Figure 5**). GFHK99 virus was examined in DF-1 and MDCK cells and MaMN99 virus was tested in MDCK cells. To ensure viral genomes were supplied as low or single copies, a dose of 0.5 RNA copies per cell was used for low MOI infection. Under these low MOI conditions in MDCK cells, GFHK99 genomic RNA levels remain low throughout the time course. GFHK99 mRNA increases early, perhaps reflecting robust primary transcription, then accumulates at a relatively slow rate from 2 h post-infection onward (**Figure 5A**). Both viral RNA species accumulate at a significantly higher rate for GFHK99 virus in DF-1 cells and MaMN99 virus in MDCK cells (**Figure 5B, C, G**). For the high MOI dose, we used 3.0 HA-expressing units per cell, as determined by flow cytometry (**Supplementary Figure 3**). This measure of infectivity gives a functional readout for polymerase activity, and therefore ensures the vast majority of cells carry an active viral polymerase. At this high MOI, accumulation of GFHK99 mRNA and vRNA in MDCK cells occurs at a similar rate to that seen for GFHK99 virus in DF-1 cells or MaMN99 virus in MDCK cells (**Figure 5D–F, H**). Thus, a host-specific defect in GFHK99 polymerase activity that affects both replication and transcription is seen at low MOI. This defect is, however, resolved under conditions where multiple infection is prevalent.

**Figure 5.**
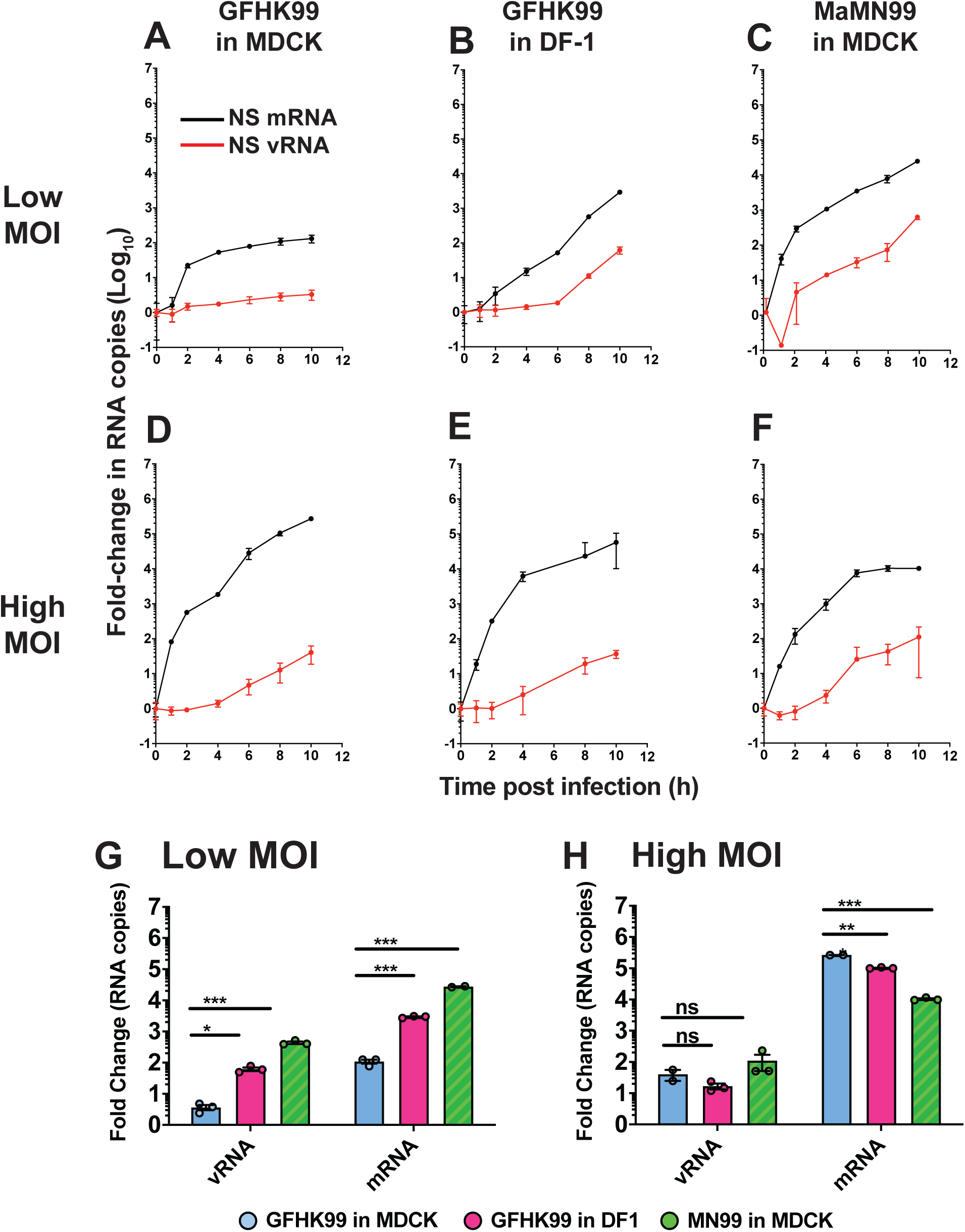
High multiplicity of infection is needed for robust GFHK99 polymerase activity in MDCK cells. Dishes of MDCK or DF-1 cells (n=3) were infected with GFHK99 or MaMN99 virus at low (0.5 RNA copies per cell) or high (3 HA-expressing units per cell) MOI. NS segment vRNA and mRNA was quantified at the indicated time points (A–F). The average fold change from initial (t=0) to peak RNA copy number is plotted for low MOI infections (G) and high MOI infections (H). Error bars represent standard deviation. Significance was assessed by multiple unpaired t-tests with correction for multiple comparisons using the Holm-Sidak method, with alpha = 5.0%. Each row was analyzed individually, without assuming a consistent SD: *p < 0.05, **<0.01, ***<0.001. ns = not significant.

### Single-cell mRNA sequencing reveals a need for cooperation

To evaluate the heterogeneity of viral RNA synthesis at the single-cell level, we infected DF-1 or MDCK cells with GFHK99 virus under single-cycle conditions and collected cells at 8 h post-infection for mRNA barcoding on the 10x Genomics Chromium platform prior to sequencing. Cells in which no viral mRNA was detected were excluded from further analysis. In addition, we used a thresholding approach to exclude cells that were unlikely to be truly infected, but rather may carry small amounts of viral mRNA owing to lysis of neighboring cells (**Supplementary Figure 4**). Following these two exclusion steps, the number of cells remaining ranged between 110 and 440 per infection condition. Next, to allow comparison between cell types and between experiments, each cell’s total transcript abundance was normalized to the median number of transcripts per cell in the relevant sample. The normalized and log_10_ transformed counts of viral transcripts per cell were then compared across conditions. We found that the amount of detected GFHK99 viral mRNA varies widely between individual DF-1 cells, which is consistent with previous observations^38–40^. In contrast, GFHK99 viral mRNA levels are uniformly low in MDCK cells, especially at the lower MOIs tested (**Figure 6A**, left facet).

**Figure 6.**
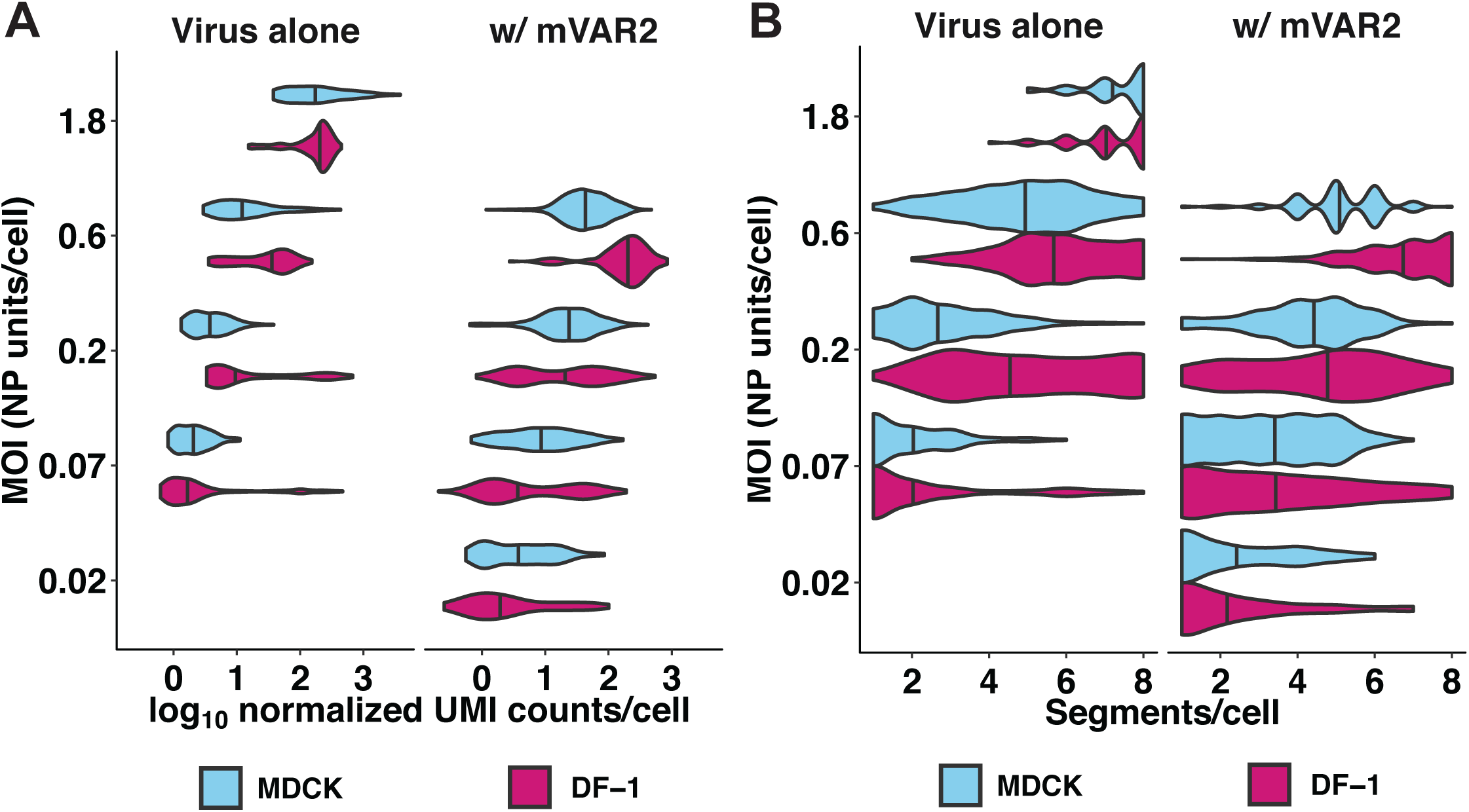
Homologous coinfecting virus boosts GFHK99 viral transcription in single cells and reveals comparable rates of segment detection in MDCK and DF-1 cells. DF-1 or MDCK cells were infected with GFHK99 virus (left facet) at MOIs of 0.07, 0.2, 0.6, or 1.8 NP-expressing units per cell, or a 1:1 mixture of GFHK99 WT and GFHK99 mVAR_1_ viruses at four different total MOIs (0.02, 0.07, 0.2, 0.6 NP-expressing units per cell) and a constant amount of GFHK99 mVAR_2_ virus (0.1 PFU per cell in DF-1 cells, 1.0 PFU per cell in MDCK cells) (right facet). Violin plots show the distribution of total viral RNA (A) or distinct viral genome segments (B) per cell in each cell-MOI infection condition. Vertical lines denote the median of each distribution. UMI = unique molecular identifier.

To evaluate whether low transcript abundance corresponded with the failure to detect one or more polymerase-encoding segments, we stratified the data based on detection of all four segments necessary to support transcription (PB2, PB1, PA and NP). Consistent with prior reports^38^, viral transcript levels are higher in cells that contained PB2, PB1, PA, and NP transcripts compared to those in which one or more of these transcripts was not detected (**Supplementary Figure 5**). We noted that the PB2, PB1, and PA mRNAs were markedly less abundant than the other viral transcripts, however. Thus, the correspondence between failure to detect polymerase transcripts and low viral mRNA abundance may simply be due to the higher likelihood of these mRNAs falling below the limit of detection in cells that have low overall viral transcription.

To measure the impact of coinfecting viruses in individual cells, we repeated the single-cell sequencing experiment with the addition of two variants of GFHK99 virus, GFHK99 mVAR_1_ and GFHK99 mVAR_2._ Cells were inoculated with GFHK99 WT and GFHK99 mVAR_1_ viruses in a 1:1 ratio. Coinfection with GFHK99 mVAR_2_ virus was performed simultaneously and this virus was used at the concentration found to give optimal support of WT viral RNA replication in **Figure 3D**: 0.1 PFU per cell in DF-1 cells and 1.0 PFU per cell in MDCK cells. After mRNA sequencing, only cells in which transcripts from all eight mVAR_2_ virus segments were detected were analyzed further. Cells that likely obtained WT or mVAR_1_ viral RNA through lysis were excluded (**Supplementary Figure 4**). Between 64 and 217 cells per infection condition met both criteria to be included in further analysis. As expected, no significant difference was detected between WT and mVar_1_ transcript levels (p > 0.05, linear mixed effects model) (**Supplementary Figure 5**). No differentiation is therefore made between strains in further analysis. In cells infected with both WT and mVAR_1_ viruses, viral transcript abundance is calculated separately for each, so that coinfected cells contribute two distinct measurements to the dataset. The viral transcript levels per cell detected in this second experiment are shown in **Figure 6A** alongside data from the first experiment. In comparing the two infections, we observe that total viral transcript abundance is markedly lower in MDCK cells compared to DF-1 cells in the first infection, but this effect is almost entirely mitigated by the presence of mVAR_2_ virus in the second infection. This reduction in the disparity between DF-1 and MDCK cells resulted from the fact that mVAR_2_ virus increased transcript abundance by 149% in DF-1 cells, but 258% in MDCK cells (p < 10^−4^, linear mixed effects model) (**Figure 6A**). These data underscore the significance of viral collective interaction to ensure productive infection in diverse hosts.

Because only a subset of a cell’s transcripts is captured and therefore reliably detected^41^, the 10x platform does not allow a robust determination of segment presence or absence in a cell. The addition of mVAR_2_ virus improves the sensitivity with which WT transcripts can be detected, however, and – as noted above – levels the playing field between MDCK and DF-1 cells. We therefore evaluated the number of distinct viral gene segments detected per cell, as indicated by the presence of the corresponding mRNAs. Failure to detect a subset of the eight segments in a given cell is very common in our datasets, as with others^38, 40^. The addition of mVAR_2_ virus increases the number of segments per cell detected, however, suggesting that failure to detect is often a result of transcript levels falling below the limit of detection of the assay (**Figure 6B**). Importantly, in the presence of mVAR_2_ virus, the number of segments detected per cell was comparable between MDCK and DF-1 cells across all MOIs tested (**Figure 6B**). This result suggests that the frequency with which incomplete GFHK99 viral genomes are expressed is comparable between these mammalian and avian cell lines.

### Frequency of incomplete GFHK99 genomes in MDCK cells is moderate

Given the limitations of single-cell mRNA sequencing for the detection of relatively rare transcripts, we employed a second single-cell approach to quantify the frequency with which fewer than eight vRNAs are replicated in GFHK99 infected MDCK cells^19^. MDCK cells were coinfected with a low MOI of GFHK99 WT virus and a high MOI of GFHK99 VAR_2_ virus. The GFHK99 VAR_2_ virus acts to ensure propagation of the WT virus gene segments, even when less than the full WT viral genome is available for transcription and replication. Following inoculation, single cells were sorted into wells which contain a naïve cell monolayer and multicycle replication was allowed for 48 h. To determine which viral gene segments were present in the initially sorted cell, RT-qPCR with primers that differentiate WT and VAR_2_ gene segments was applied. As detailed in the Methods, the results were used to calculate the probability that a cell infected with a single WT virus contains a given segment. We termed the resultant parameter Probability Present (*P_P_*). The experimentally determined *P_P_* values vary among the segments, with a range of 0.56 to 0.8 (**Figure 7A**). The product of the eight *P_P_* values gives an estimate of the proportion of singular infections in which all eight segments are available for replication. This estimate is 3.2% for GFHK99 virus in MDCK cells.

**Figure 7.**
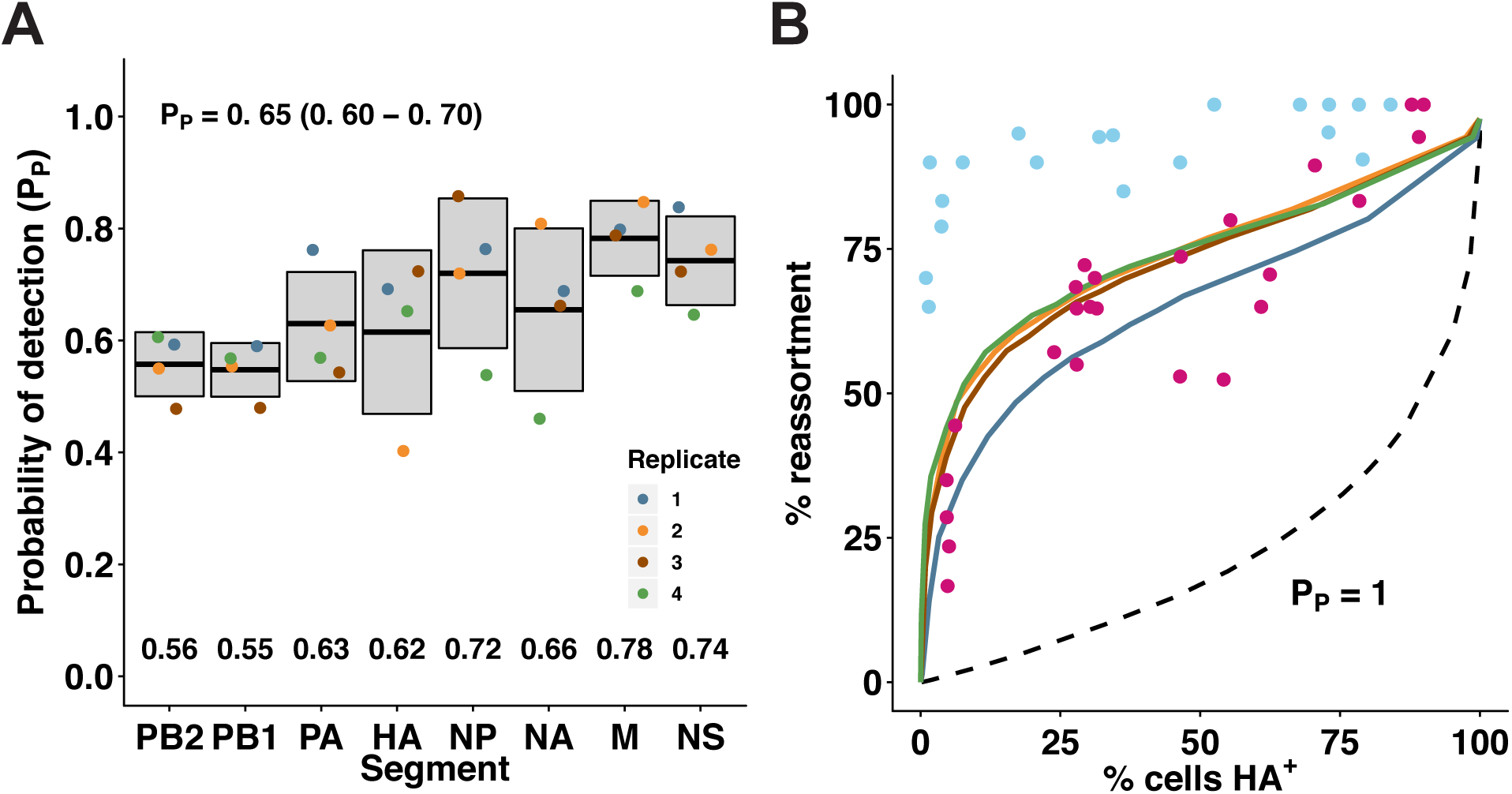
Incomplete GFHK99 virus genomes are present but not sufficiently abundant to account for observed reassortment in MDCK cells. Incomplete viral genomes were quantified experimentally by a single-cell assay which relies on the amplification of incomplete viral genomes of GFHK99 WT virus (0.018 PFU per cell) by a genetically similar coinfecting virus, GFHK99 VAR_2_. Based on the rate of detection of GFHK99 WT virus segments in this assay, the probability that a given segment would be present and replicated in a singly infected MDCK cell is reported as *P_P_*. A) Summary of experimental *P_P_* data. n = 4 biological replicates, distinguished by color. Horizontal bars and shading represent mean ± standard deviation. Mean *P_P,i_* values for each segment are indicated at the bottom of the plot area. Average *P_P_* for each experiment was calculated as the geometric mean of the eight *P_P,i_* values, and the mean ± standard deviation of those four summary *P_P_* values is shown at the top of the plot area. B) Experimentally obtained *P_P,i_* values in A were used to parameterize a computational model^18^. Levels of reassortment predicted using the experimentally determined parameters are shown with colors corresponding to points in (A). Levels of reassortment predicted if *P_P_*=1.0 are shown with the dashed line. Observed reassortment of GFHK99 WT and VAR viruses in MDCK cells are shown with blue circles. Observed reassortment of GFHK99 WT and VAR viruses in DF-1 cells are shown with pink circles. Observed data are the same as those plotted in Figure 1.

To evaluate whether incomplete viral genomes account for the focusing of GFHK99 virus production within multiply infected cells, we used our previously published computational model of IAV coinfection and reassortment^18^. In this model, the frequency of successful segment replication is governed by eight *P_P_* parameters and an infected cell only produces virus if at least one copy of all eight segments are replicated. Importantly, in this model the amount of virus produced from productively infected cells is constant – there is no additional benefit to multiple infection. When the eight experimentally determined *P_P_* values for GFHK99 virus in MDCK cells are used to parameterize the model, the theoretical prediction of reassortment frequency is much lower than that observed experimentally for GFHK99 viruses in MDCK cells, but a good match with the reassortment seen in DF-1 cells (**Figure 7B**). Clearly, the frequency with which GFHK99 segments fail to be replicated cannot fully account for the high reassortment seen in the mammalian MDCK cell system, but can fully account for the more moderate reliance on multiple infection seen in DF-1 cells. Thus, in mammalian cells, a full GFHK99 viral genome is necessary but not sufficient to support robust replication.

## Discussion

Our data reveal that moderate reliance of IAV on multiple infection is the norm. An exceptionally high need for multiple infection can, however, occur when an IAV infects a new host species. Thus, our findings identify increased reliance on multiple infection as a novel component of IAV host restriction. Since not all mismatched virus-host combinations show a high reliance on multiple infection, the data indicate that a need for collective interactions is a potential manifestation of IAV host restriction, not a universal feature. Dependence on multiple infection is of particular interest as a barrier to cross-species transfer for two reasons: first, it can be overcome in the absence of genetic adaptation, through high dose infection; and second, it leads to high levels of reassortment, which in turn can facilitate adaptation to a new host.

An important consequence of viral genome segmentation is the potential for replication of incomplete genomes^2^. Complementation is therefore a major class of collective interaction for viruses with segmented genomes. The relevance of complementation for a given virus species likely depends on the extent to which packaging, delivery and replication of genome segments is coordinated^42^. For IAV, we and others have demonstrated that a subset of segments fails to be replicated or expressed in infected cells with high frequency^16, 19, 38, 40^. Specifically, for influenza A/Panama/2007/99 (H3N2) virus in MDCK cells, we found that delivery of a single viral genome results in replication of all eight segments only 1.2% of the time^19^. Data reported herein for GFHK99 virus indicate that a somewhat higher proportion of replicated viral genomes are complete – namely, 3.2%. Thus, GFHK99 virus is partially dependent on complementation for productive infection. However, the high levels of reassortment seen between GFHK99 WT and VAR viruses in mammalian cells indicate that additional cooperative interactions are at play. This is clear from the discrepancy between observed GFHK99 virus reassortment in MDCK cells and the reassortment levels expected if complementation is the only cooperative effect considered. These data point to a model in which the presence of not just complete genomes, but rather multiple copies of the viral genome, are needed to overcome host-specific barriers to GFHK99 infection in mammalian systems (**Figure 8**).

**Figure 8.**
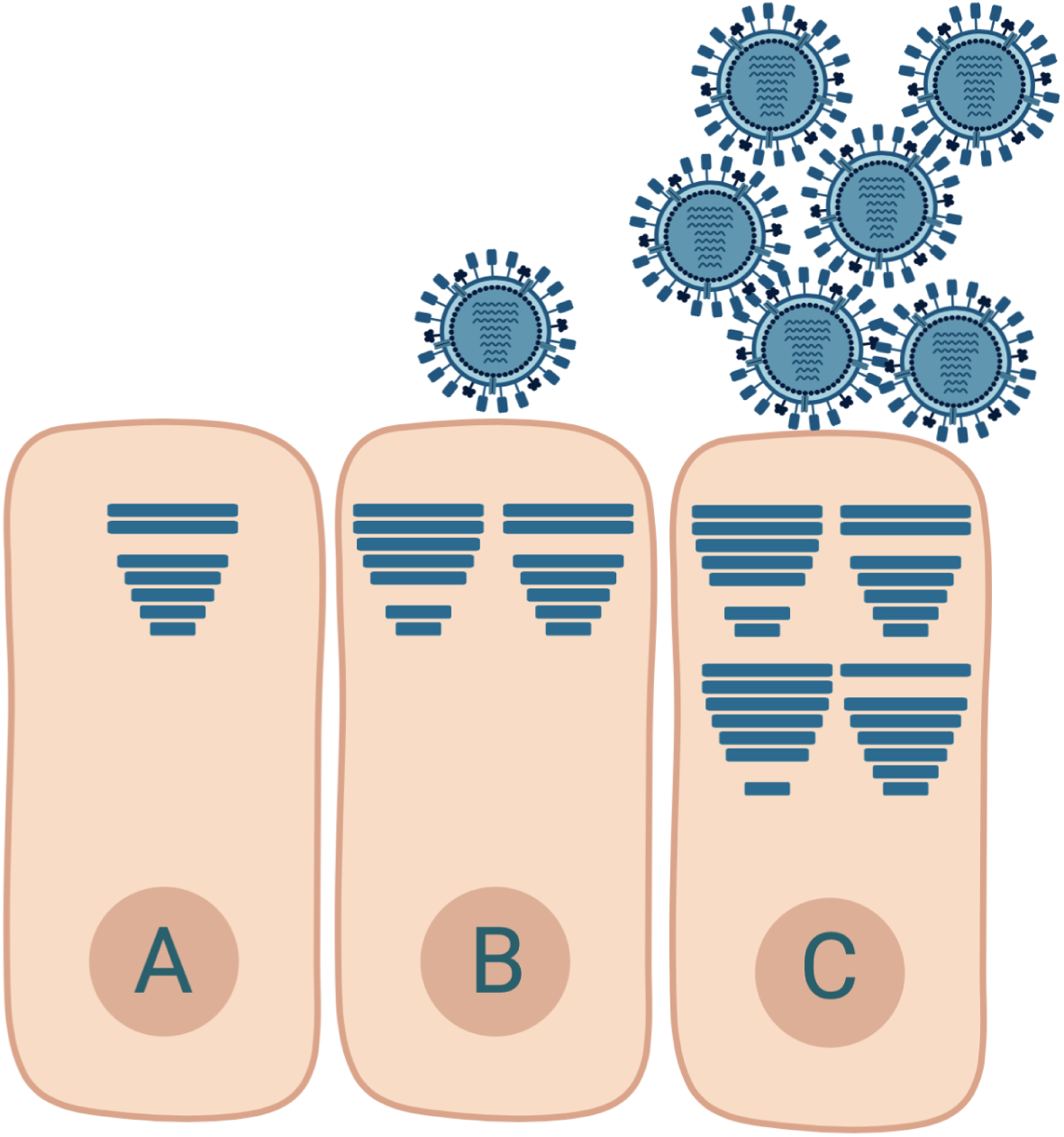
Model: complementation of incomplete viral genomes is necessary but not sufficient for robust progeny production from mammalian cells infected with GFHK99 virus. Cells infected with single virions often replicate incomplete viral genomes and therefore do not produce viral progeny (A). In mammalian cells, complementation of incomplete viral genomes through coinfection may allow progeny production (B), but robust viral yields require the delivery of additional genomes to the cell, beyond the levels needed for complementation (C). Horizontal bars within cells represent viral gene segments successfully delivered and replicated.

Insight into the nature of this second cooperative interaction is gleaned from the observation that the amount of viral RNA produced from GFHK99 WT viral genomes is significantly increased with the addition of a homologous virus. From these data, we can conclude that the coinfecting virus functions in *trans* to support WT virus RNA replication in mammalian cells. In addition, our genetic data implicate the viral PA in defining an acute reliance on cooperation. The PA gene segment encodes two proteins: PA and PA-X^43^. Together with PB2 and PB1, PA is an essential component of the trimeric polymerase complex^44^. PA carries the endonuclease activity required for cap-snatching from nascent Pol II transcripts, which is essential for viral transcription^45^. It also comprises much of the interface for viral polymerase dimerization, which is needed for viral replication^46^. PA-X shares its N-terminus with PA but has a unique 41 or 61 amino acid C-terminus^43, 47^. PA-X contributes to the shut-off of host protein synthesis by targeting host Pol II transcripts for destruction^48, 49^. Alignment of PA and PA-X proteins encoded by GFHK99 and MaMN99 viruses reveals 29 differences in PA and 9 differences in PA-X (**Supplementary Figure 6**). Three-dimensional modeling of the GFHK99 and MaMN99 PA proteins based on the recently reported structure of the A/NT/60/1968 (H3N2) virus polymerase^46^ did not predict marked differences. Nevertheless, the polymorphisms noted in and near the dimerization loop suggest a model in which the reliance of GFHK99 virus on multiple infection may result from an inefficiency of polymerase dimerization and consequent inefficiency of vRNA synthesis^46^. This model is attractive based on the results of Fan et al., who reported that aberrant *in vitro* initiation of vRNA synthesis by a dimerization mutant was ameliorated with increased polymerase concentration^46^. We speculate that multiple infection could lead to a similar effect within the infected cell. If this speculation is correct, it would suggest a role for host factors in viral polymerase dimerization, since the observed GFHK99 reliance on multiple infection is specific to mammalian cells. Finer mapping of the phenotype within PA and targeted functional studies are needed to test this model and explore other possible mechanisms.

Of note, high reliance on multiple infection in mammalian cells does not extend to all avian IAVs. Our experiments indicate that this phenotype is a feature of H9N2 subtype avian IAV prevalent at the poultry-human interface, while two avian IAV derived from wild birds, MaMN99 and DkHK78 viruses, showed no increase or a modest increase in reliance on multiple infection in mammalian relative to avian systems. Whether a need for multiple infection in mammalian cells extends to other poultry-adapted lineages is an interesting question for future study. H9N2 subtype viruses are, however, highly relevant in the context of zoonotic infection due to their prevalence at the poultry-human interface. Sporadic human infections with poultry-adapted H9N2 viruses have been reported, including with strains closely related to GFHK99 virus. These H9N2 viruses furthermore share several related genes with H5N1 and H7N9 subtype viruses that have caused hundreds of severe human infections^50–55^. The G1 lineage to which the GFHK99 virus belongs circulates widely in the poultry of Southeast Asia, the Middle East, and North Africa and reassorts frequently with other poultry adapted IAVs^56–59^. Our comparison of reassortment in guinea pigs and quail indicates that reassortment could be particularly prevalent in the context of zoonotic infection of mammals. This phenotype of high reassortment in mammals is expected to extend to the H5N1 and H7N9 subtype viruses of public health concern, which carry polymerase genes related to those of the GFHK99 virus^53–55^. While reassortment is typically deleterious^60, 61^, high frequencies increase opportunity for fit genotypes to arise and adapt, and should therefore be considered in assessing the risk of emergence posed by non-human adapted IAVs.

Our data reveal novel concepts in both virus-host and virus-virus interactions. In the context of virus-host interactions, the extent of IAV reliance on cooperation is determined in part by interactions with the host cell, such that infection of novel hosts may be characterized by an acute dependence on multiple infection. In the context of virus-virus interactions, our data show that cooperation is not limited to complementation; rather, interactions among homologous coinfecting viruses can be highly biologically significant. Viral dependence on multiple infection is expected to have a strong impact on viral fitness and evolution by modulating the efficiency of spread within and between hosts and by changing how a virus population samples sequence space^62–65^. Thus, multiple infection dependence is likely to play an important role in determining the outcomes of IAV infection and evolution in diverse hosts.

## Acknowledgements

We thank David Stallknecht (University of Georgia) for providing the A/mallard/MN/199106/99 (H3N8) biological isolate. We thank Hui Tao, Shamika Danzy, and Ginger Geiger for technical assistance. This work was funded in part by NIH R01 AI127799 (to ACL and DRP); NIH/NIAID Centers of Excellence in Influenza Research and Surveillance (CEIRS), contract numbers HHSN272201400004C (to ACL and GST) and HHSN272201400008C (to DRP). Additional funds were provided by the Georgia Research Alliance and the Georgia Poultry Federation (to DRP) and NIH/NIAID Genomic Centers for Infectious Diseases (GCID), Award Number U19AI110819.

## Author Contributions

KLP contributed to the conception of work, experimental design, data acquisition and analysis, interpretation of data; KG, NTJ and C-YL contributed to experimental design, data acquisition and analysis and interpretation of data; SC contributed to data acquisition; MM and MCW contributed to data acquisition and analysis; BEP contributed to data analysis and interpretation; GST contributed to conception of the work, data analysis and interpretation; LMF contributed to data acquisition; DRP contributed to experimental design, data analysis and interpretation; ACL contributed to conception of work, experimental design and data analysis and interpretation. All authors contributed to the writing of the manuscript.

## Data availability

10x Genomics single-cell sequencing data is available on the GEO database with the accession number XXXXXXXX. Other data are available from the corresponding author upon reasonable request.

## Code availability

Code used for 10x Genomics analysis is available at: https://github.com/njacobs627/GFHK99_Multiplicity. Code used to run the agent-based model on influenza A virus reassortment was reported previously^18, 19^ and is also available at https://github.com/njacobs627/GFHK99_Multiplicity.

## Methods

### Cells and cell culture media

Madin-Darby canine kidney (MDCK) cells, a gift from Peter Palese, Icahn School of Medicine at Mount Sinai, were used in all experiments. MDCK cells from Daniel Perez at University of Georgia were used for plaque assays as this variant of the MDCK line was found to yield more distinct plaques for the GFHK99 strain. Both MDCK cell lines were maintained in minimal essential medium (MEM; Gibco) supplemented with 10% fetal bovine serum (FBS; Atlanta Biologicals), penicillin (100 IU), and streptomycin (100 µg per mL) (PS; Corning). 293T cells (ATCC CRL-3216) and DF-1 cells (ATCC CRL-12203) were maintained in Dulbecco’s minimal essential medium (DMEM; Gibco) supplemented with 10% FBS and PS. Duck embryo cells (ATCC CCL-141) were maintained in Eagle’s minimum essential medium (EMEM; ATCC) supplemented with 10% FBS and PS. Normal human bronchial epithelial (NHBE) cells were acquired from Lonza and were amplified and differentiated into air-liquid interface cultures as recommended by Lonza and described by Danzy et al.^66^. All cells were cultured at 37°C and 5% CO_2_ in a humidified incubator. Medium for culture of IAV in each cell line (virus medium) was prepared by supplementing the appropriate media with 4.3% bovine serum albumin and penicillin (100 IU), and streptomycin (100 µg per mL). Ammonium chloride-containing virus medium was prepared by the addition of HEPES buffer and NH_4_Cl at final concentrations of 50 mM and 20 mM, respectively.

### Viruses

All viruses were generated by reverse genetics^67^. For avian viruses, 293T cells transfected with reverse genetics plasmids 16–24 h prior were injected into the allantoic cavity of 9–11 day old embryonated chicken eggs and incubated at 37°C for 40–48 h. The resultant egg passage 1 stocks were used in experiments. For NL09-based viruses, 293T cells transfected with reverse genetics plasmids 16–24 h prior were co-cultured with MDCK cells at 37°C for 40–48 h. Supernatants were then propagated in MDCK cells from low MOI to generate NL09 working stocks. Defective interfering segment content of PB2, PB1, and PA segments was confirmed to be minimal for each virus stock, as described previously^68^ (**Supplementary Figure 7**). All plaque assays were performed in MDCK cells; viruses were also titered by flow cytometry and/or RT qPCR-based methods where indicated. The reverse genetics system for influenza A/guinea fowl/Hong Kong/WF10/99 (H9N2) virus was reported previously^69, 70^. This strain has been referred to as WF10 in previous publications^69–71^. For consistency with other strains used in the present manuscript, it is referred to herein as GFHK99. A low passage isolate of influenza A/mallard/Minnesota/199106/99 (H3N8) virus, referred to herein as MaMN99, was obtained from David Stallknecht at the University of Georgia^72^. The virus was passaged once in eggs and then the eight cDNAs were generated and cloned into the pDP2002 vector^73^. A low passage isolates of influenza A/duck/Hong Kong/448/1978 (H9N2) and A/quail/Hong Kong/A28945/1988 (H9N2), referred to herein as dkHK78 and QaHK88, were passaged once in eggs and then the eight cDNAs were generated and cloned into the pDP2002 vector. To increase the recovery efficiency of viruses containing polymerase components from the MaMN99 virus, pCAGGS support plasmids encoding PB2, PB1, PA, and NP proteins of the A/WSN/33 (H1N1) strain were supplied. GFHK99 WT, MaMN99 VAR and NL09 VAR viruses were engineered to contain a 6XHis epitope tag plus GGGS linker at the N-terminus of the HA protein following the signal peptide.

GFHK99 VAR_1_, MaMN99 WT and NL09 VAR viruses contain similarly modified HA genes, with an HA epitope tag plus a GGGS linker inserted at the N-terminus of the HA protein^36^. All assays except the single-cell mRNA sequencing used viruses carrying these epitope tags.

Silent mutations were introduced into VAR viruses by site-directed mutagenesis to allow genotyping of WT and VAR segment origin. The specific changes introduced into all VAR viruses used here are listed in **Supplementary Table 1**. Mutations introduced into GFHK99 VAR_1_, MaMN99 VAR, and NL09 VAR viruses enable detection by high-resolution melt analysis or probe-based droplet digital PCR. Mutations introduced into the GFHK99 VAR_2_ strain were designed to confer unique primer binding sites relative to GFHK99 WT virus. Viruses used for single-cell mRNA sequencing, GFHK99 mVAR_1_ and GFHK99 mVAR_2_, carry mutations near the 3’ end of each transcript to allow detection by sequencing following oligo-dT priming^38^.

### Synchronized, single-cycle infection conditions

Conditions designed to synchronize viral entry and prevent propagation of progeny viruses were used for all cell culture-based infections with the exception of those performed for measurement of *P_P_* values. Synchronized, single-cycle infections were performed as follows: Cell monolayers were washed three times with PBS and placed on ice. Chilled virus inoculum was added to each well and incubated at 4°C for 45 minutes with occasional rocking to allow attachment without entry^74^. Inoculum was aspirated and cell monolayer was washed three times with cold PBS before addition of warm virus medium lacking trypsin. Cultures were incubated at 37°C. At 3 h post-infection, virus medium was replaced with ammonium chloride-containing virus medium. Addition of ammonium chloride to the medium prevents acidification of endosomes, thereby blocking further IAV infection^75^. Cultures were returned to 37°C for the remainder of the incubation time.

### Infection of cultured cells for quantification of coinfection and reassortment

MDCK or DF-1 cells were seeded at a density of 4×10^5^ cells per well in 6-well dishes 24 h before inoculation. Virus inoculum was prepared by combining WT and VAR viruses at high titer in a 1:1 ratio based on PFU titers, and then diluting in PBS to achieve MOIs ranging from 10 to 0.01 PFU per cell. Synchronized, single cycle infection conditions were used. In addition, owing to the low yield of IAV in DF-1 cells, acid inactivation of inoculum virus was performed at 1 h post-infection for this cell type. This procedure is needed if residual inoculum virus would otherwise comprise an appreciable proportion of the virus sampled at 12 or 16 h post-inoculation. For acid inactivation, media was aspirated and replaced with 500 µL of PBS-HCl, pH 3.00 and incubated 5 min at 37°C. Cells were then washed once with PBS before the addition of virus medium. GFHK99 virus-infected cells were harvested at 12 h post-infection due to high amounts of cytopathic effects at later time points. Cells infected with MaMN99 virus and MaMN99:GFHK99 chimeric viruses were harvested at 16 h post-infection. NL09 virus reassortment data shown in Figure 1C were reported previously^76^ and are included here to allow comparison to the avian strains used.

### Determination of infection and coinfection levels based on HA surface expression

To enumerate infected cells, surface expression of HIS and HA epitope tags was detected by flow cytometry (**Supplementary Figure 8**). This method was previously described in detail^36^. Cells were fixed following staining by resuspending in FACS buffer and 1% paraformaldehyde. The percentage of cells positive for either or both epitope tags is expressed as % cells HA^+^. The percentage of cells positive for both epitope tags is expressed as % cells dual-HA^+^. The relationship between these two parameters was evaluated by plotting % cells dual-HA^+^ against % cells HA^+^ and regressing the resultant curve as a polynomial:

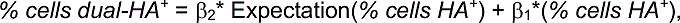

where β_2_ and β_1_ are genotype-specific. For a given fraction of cells expressing HA, the expected fraction of cells expressing both HA proteins is derived from the Poisson distribution with λ = –*ln*(1 – *% cells HA*^+^):

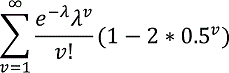

From the regression models, we then quantified the degree of linearity using the equation:

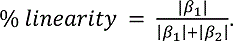

### Animal models and reassortment *in vivo*

Ethics Statement. All animal experiments were conducted in accordance with the Guide for the Care and Use of Laboratory Animals of the National Institutes of Health. Studies were conducted under animal biosafety level 2 (ABSL-2) containment and approved by the IACUC of the University of Georgia (protocols A201506-026-Y3-A2 and A201506-026-Y3-A5) for quail studies or the IACUC of Emory University (protocol PROTO201700595) for guinea pig studies. Animals were humanely euthanized following guidelines approved by the American Veterinary Medical Association (AVMA).

Quail eggs obtained from the College of Veterinary Medicine, University of Georgia, were hatched at the Poultry Diagnostic and Research Center, University of Georgia. Two days before virus inoculation, quail sera were confirmed to be seronegative for IAV exposure by NP ELISA (IDEXX, Westbrook, ME). At 3 weeks of age, birds were moved into a HEPA in/out BSL2 facility and each group divided into individual isolator units.

Groups (n=6) of 3 week old Japanese quail (Coturnix Japonica) were used to determine the 50% quail infectious dose of the 1:1 GFHK99 WT and GFHK99 VAR_1_ virus mixture. Each quail was inoculated with 500 µl by oculo-naso-tracheal route of virus mixture in PBS, at increasing concentrations of 10^0^ to 10^6^ TCID_50_ per 500 µL. Tracheal and cloacal swab specimens were collected daily from each bird in brain heart infusion media (BHI). Swab samples were analyzed by TCID_50_ assay and titers of tracheal swabs collected at 4 d post-inoculation were used to determine the QID_50_ by the Reed and Muench method^77^. Virus was not detected in cloacal swabs. QID_50_ was found to be equivalent to 1 TCID_50_.

To quantify reassortment in quail, samples collected from quail (n=6) infected with the 10^2^ TCID_50_ dose of the 1:1 GFHK99 WT and GFHK99 VAR_1_ virus mixture were used. These were the same birds as used to determine QID_50_. Virus shedding kinetics were determined by plaque assay of tracheal swab samples and samples from days 1, 3, and 5 were chosen for genotyping of virus isolates.

Female Hartley strain guinea pigs weighing 250–350 g were obtained from Charles River Laboratories (Wilmington, MA) and housed by Emory University Department of Animal Resources. Prior to intranasal inoculation and nasal washing, guinea pigs were sedated by intramuscular injection with 30 mg per kg ketamine/ 4 mg per kg xylazine. The GPID_50_ of GFHK99 WT/VAR_1_ and MaMN99 WT/VAR virus mixtures were determined as follows. Groups of four guinea pigs were inoculated intranasally with virus mixture in PBS at doses of 10^0^ to 10^5^ PFU per 300 µL inoculum. Daily nasal washes were collected in 1 mL PBS and titered by plaque assay. Results from day 2 nasal washes were used to determine the GPID_50_ by the Reed and Muench method^77^. The GPID_50_ of GFHK99 virus was found to be 2.1 x 10^3^ PFU; that of MaMN99 virus was 2.1 x 10^1^ PFU. The GPID50 of NL09 virus was previously determined to be 1 x 10^1^ PFU (ref^78^ and data not shown).

To evaluate reassortment kinetics in guinea pigs, groups of six animals were infected with 10^2^ x GPID_50_ of the aforementioned GFHK99 WT / VAR_1_, MaMN99 WT / VAR or NL09 WT / VAR virus mixtures. Virus inoculum was given intrasnasally in a 300 µL volume of PBS. Nasal washes were performed on days 1–6 post-inoculation and titered for viral shedding by plaque assay. HRM genotyping was performed on samples collected on day 1, 3, and 5 for each guinea pig.

### Quantification of reassortment and effective diversity

Reassortment was quantified for *in vitro* coinfection supernatants, guinea pig nasal washes, and quail tracheal swabs as described previously^36^. Briefly, plaque assays were performed in 10 cm dishes to isolate virus clones. 1 mL serological pipettes were used to collect agar plugs into 160 µl PBS. Using a ZR-96 viral RNA kit (Zymo), RNA was extracted from the agar plugs and eluted in 40 µl nuclease free water (Invitrogen). Reverse transcription was performed using Maxima RT (Thermofisher) according to the manufacturer’s protocol. The resulting cDNA was diluted 1:4 in nuclease free water and each cDNA was combined with segment specific primers (**Supplementary Table 2**) and Precision Melt Supermix (Bio-Rad) and analyzed by qPCR in a CFX384 Touch real-time PCR detection system (Bio-Rad) designed to amplify a ∼100 bp region of each gene segment which contains a single nucleotide change in the VAR virus. The qPCR was followed by high-resolution melt (HRM) analysis to differentiate WT and VAR amplicons^79^. Precision Melt Analysis software (Bio-Rad) was used to determine the parental virus origin of each gene segment based on melting properties of the cDNAs and comparison to WT and VAR controls. Each individual plaque was assigned a genotype based on the combination of WT and VAR genome segments, with 2 variants on each of 8 segments allowing for 256 potential genotypes.

Reassortment data was used to calculate viral genotypic diversity; that is, the diversity of the 256 possible WT/VAR genotypes present in a given sample. Diversity was quantified as reported previously^80^, by calculating Simpson’s Index, given by *D* = sum(*p_i_*^2^), where *p_i_* represents the proportional abundance of each genotype^81^. Simpson’s Index accounts for both the raw number of species and variation in abundance of each, and is sensitive to the abundance of dominant species. These features are important to ensure that overrepresentation of a single reassortant genotype does not cause a viral population to appear more diverse than it actually is. Because Simpson’s Index does not scale linearly, each sample’s Simpson’s Index value was converted to a corresponding Hill number to derive its effective diversity, *N*_2_ = 1/*D*^82^, which is defined as the number of equally abundant species required to generate the observed diversity in a sample community. Because it scales linearly, Hill’s *N*_2_ allows a more intuitive comparison between communities (i.e., a community with *N*_2_ = 10 species is twice as diverse as one with *N*_2_ = 5) and is suitable for statistical analysis by basic linear regression methods^83^. Robust linear models of *N_2_* vs. time were regressed using the R package *robustlmm*.

### Hierarchical clustering analysis

To compare the behavior of multiple viruses during *in vitro* coinfections, a hierarchical clustering algorithm was used. In these experiments, multiplicity dependence was measured by analyzing HA co-expression and reassortment as a function of the percentage of cells expressing HA. Each of these regression models contain two parameters (*% cells dual-HA^+^* = β_2_* Expectation(*% cells HA^+^*) + β_1_*(*% cells HA^+^*), and *% reassortment* = β_0_ + β_1_ * *ln*(*% cells HA^+^*). The behavior of a given virus can therefore be expressed as a set of four parameters. These parameters were used to calculate the distance between each pair of viruses, and Ward’s method of aggolomerative hierarchial clustering was then used to organize viruses into clusters based on these distances^84^. Briefly, this algorithm combines a nearby pair of elements into a cluster with a new position that is halfway between the original individuals. This process is repeated until all elements have been incorporated into one cluster. The R package *pvclust* was used to calculate statistical support for the existence of each node by multiscale bootstrap resampling, with nodes appearing in 95% of trials being deemed statistically significant. Each dendrogram was then divided into two clusters to determine whether the behavior of GFHK99 virus in MDCK cells represented a true outgroup (**Figure 1**), or to determine whether each GFHK99:MaMN99 chimeric virus was more similar to GFHK99 or MaMN99 parent virus (**Figure 4**).

### Single-cycle viral growth kinetics

DF-1 or MDCK cells were seeded at 4×10^5^ cells per well in 6 well dishes 24 h prior to infection. Virus was serially diluted using PBS. Synchronized, single cycle infection conditions with acid inactivation of inoculum virus as described above were used. At each time point, 120 µl supernatant was collected. Viral titers for each sample were assessed by plaque assay in MDCK cells. Each MOI condition was used in 5–6 wells in parallel infections. Three wells served as technical replicates for growth curve sampling while the remaining wells were harvested at 24 h post-infection to enumerate HA-expressing cells via flow cytometry. Flow cytometry data for NL09 in MDCK cells are not included owing to low sensitivity of the assay for this virus. In cases where acid inactivation was inefficient, the replicate was eliminated, and data are plotted in duplicate.

### Effect of increasing multiple infection on viral RNA replication

For DF-1 and MDCK cell experiments, 12-well plates were seeded with 3×10^5^ cells per well 24 h prior to infection. For NHBE cells, cells were cultured at an air-liquid interface as previously described^66^. Cell surfaces were washed three times with PBS prior to inoculation. Triplicate wells were then inoculated with increasing doses of VAR virus, plus 0.005 PFU per cell of WT virus in DF-1 and MDCK cells or 0.05 PFU per cell of WT virus in NHBE cells. VAR viruses used were GFHK99 VAR_2_, MaMN99 VAR, NL09 VAR, MaMN99-Anhui-PA VAR, MaMN99-GFHK99-PA VAR, dkHK78 VAR, and QaHK88 VAR. After 55 minutes at 37°C, inoculum was aspirated, cells were washed three times with PBS and 1 ml per well virus medium was added. Media was exchanged for ammonium chloride-containing media 3 h later. At 12 h post-infection, virus media was removed and cells were harvested using RNAprotect Cell Reagent (Qiagen). RNA was extracted using RNeasy columns (Qiagen) and then reverse transcribed with universal influenza primers^85^ and Maxima RT per protocol instructions. Droplet digital PCR (ddPCR) was performed on the resultant cDNA. For GFHK99 virus, QX200™ ddPCR™ EvaGreen Supermix (Bio-Rad) was used with a combination of PB2, M, and NS primers specific for the GFHK99 WT virus (final primer concentration of 200 nM) (**Supplementary Table 3**). For MaMN99, MaMN99-Anhui-PA, MaMN99-GFHK99-PA, dkHK78, QaHK88 and NL09 viruses, QX200™ ddPCR™ Supermix for Probes (Bio-Rad) was used with NP specific primers and probes (**Supplementary Table 3**).

### Strand-specific quantification of viral RNA species over time

Viruses used for this experiment were the same GFHK99 WT/VAR_1_ or MaMN99 WT/VAR virus mixtures used to measure reassortment, but in this case each mixture was considered as a single virus population (i.e. the RT ddPCR assay outlined below to quantify viral m/vRNA does not differentiate between WT and VAR genotypes). Doses used were 0.5 RNA copies per cell for the low MOI and 3.0 HA-expressing units/cell for the high MOI. RNA copy numbers of virus stocks were determined by ddPCR targeting at least four segments (mean values were used). HA-expressing units per mL was determined by counting HA^+^ cells by flow cytometry in the relevant cell type. Specifically, cells were infected with serial dilutions of virus under synchronized, single cycle conditions. At 24 h post-infection, cells were harvested and flow cytometry was performed targeting His and HA epitope tags. HA expressing units per mL for each virus / cell combination was calculated based on the linear range of %HA^+^ cells plotted as a function of volume of virus added to cells^86^ (**Supplementary Figure 3**).

12-well plates were seeded with 2×10^5^ cells per well of MDCK or DF-1 cells and incubated at 37°C for 24 h. Synchronized, single cycle infection conditions were used, as described above. Chilled virus was added at a volume of 125 µL per well. At 0, 1, 2, 4, 6, 8, and 10 h post-infection, virus medium was aspirated and cells were harvested using 400 µL of RNAprotect Cell Reagent (Qiagen). RNA was extracted using the Qiagen RNAeasy Mini kit. Two reverse transcription reactions per sample were set up with primers targeting mRNA or vRNA of the NS segment, each containing different nucleotide barcode tags (**Supplementary Table 4**). Maxima RT was used according to the manufacturer’s instructions and combined with 300 ng MDCK or 150 ng DF-1 RNA. Absolute copy number of cDNA was determined by ddPCR. Forward and reverse primers for vRNA or mRNA of NS at a total concentration of 200 nM were combined with diluted cDNA and QX200™ ddPCR™ EvaGreen Supermix (Bio-Rad). Primer sequences are given in **Supplementary Table 4**. Thermocycler protocol was 95°C for 5 min, [95°C for 30s, 57°C for 60s] repeat 40x, 4°C for 5 min, 90°C for 5 min, 4°C hold. Copy number was normalized to RNA input to give final results in units of copy number per ng RNA. We did attempt to measure cRNA in this assay but found that primers designed to be specific for cRNA cross-primed on viral mRNA.

### Single-cell mRNA sequencing

For this assay, viruses were titered in DF-1 cells using flow cytometry with anti-NP antibody. DF-1 cells were used because they give more sensitive detection of GFHK99 virus infection than MDCK cells. Cells were infected with serial dilutions of virus under synchronized, single-cycle conditions. At 24 h post-infection flow cytometry was performed targeting NP: cells were processed with the BD Cytofix/Cytoperm™ Kit (catalog no. 554714) and stained with anti-NP antibody (Abcam, clone 9G8, catalog no. ab43821) followed by donkey anti-Mouse IgG Alexa Fluor 488 (Invitrogen catalog no. A-21202). NP expression units per were calculated based on the linear range of % cells NP^+^ plotted as a function of volume of virus added to cells^86^ (**Supplementary Figure 3**).

To perform single-cell mRNA sequencing, MDCK and DF-1 cells were seeded into 6-well plates at 5×10^5^ cells per well. At 24 h post seeding, cells were infected with synchronized, single cycle conditions. In the first experiment, MDCK or DF-1 cells were inoculated with GFHK99 WT virus at an MOI of 0.07, 0.2, 0.6, or 1.8 NP units per cell. In the second experiment, MDCK or DF-1 cells were inoculated with a 1:1 ratio of GFHK99 WT virus and GFHK99 mVAR_1_ virus that amounted to a MOI of 0.02, 0.07, 0.2 or 0.6 NP units per cell. GFHK99 mVAR_2_ virus was added to MDCK and DF-1 cell infections at MOIs of 1 PFU per cell and 0.1 PFU per cell, respectively. Infected cells were incubated for 8 h at 37°C. Culture media was then aspirated and cells washed once with 1X PBS. Cells were then trypsinized with 200 µL of 0.25% Trypsin EDTA until all cells came off the plate and were mono-dispersed. To each well, 0.5 mL of virus medium was added and replicates were pooled (2 wells per MOI). Cells for each sample were counted. Samples were spun at 150 rcf for 3 minutes and washed with 0.5 mL of 1X PBS/0.04% BSA. Washings were performed two more times. Finally, cells were resuspended with 1X PBS/0.04% BSA to give 7 x 10^5^ cells per mL. Preparation for single-cell transcriptomic sequencing followed the protocol for 10x Genomics Chromium Single Cell platform.

Analysis of viral transcripts from single cells was performed with the sequencing data from all experiments using Cell Ranger software. Briefly, the Cell Ranger software assigns each read to individual cells and transcripts based on two sets of unique molecular identifiers (UMIs) that are ligated prior to amplification. This approach allows the quantification of amplification bias at both the cellular and transcript levels. The first step of the analytical workflow was to map the reads to concatenated transcriptomes of GFHK99 virus with the transcriptomes of dog or chicken to analyze MDCK and DF-1 cell infections, respectively. Protein coding regions for the dog and chicken transcriptomes were identified in the GTF file associated with genome builds CanFam3.1.98 (NCBI accession number GCA_000002285.2) and GRCg6a (NCBI accession number GCA_000002315.5), respectively, while GFHK99 virus coding regions were extracted from the reverse-complement sequences of the GFHK99 strains. The aligned sequencing data is available on the GEO database with the accession number XXXXXXXX. Cell Ranger output in the form of gene and barcode counts were then analyzed in R using the package *CellrangerRkit*. To account for the issue of cellular lysis, which can allow uninfected cells to acquire viral RNA from the supernatant and thus appear infected, a preliminary analysis was conducted to exclude cells that were likely false positives. This analysis was informed by the reasoning that 1) contaminated cells were likely to contain less viral RNA than truly infected cells, and 2) the amount of viral RNA present in contaminated cells should be less consistent than the amount present in truly infected cells. Within each infection, the proportion of each cell’s transcriptome that was comprised of viral RNA was calculated, and cells were ordered according to this proportion so that the marginal gain in % viral RNA from one cell to the next could be calculated. Excluding cells with no marginal gain (indicating two cells with the same % viral RNA), the marginal gain vs. % viral RNA was plotted for each infection, which showed that this marginal gain is initially a rapidly decreasing function of % viral RNA. The first local minimum of a local regression was derived for each infection, to determine the point at which marginal gain became more consistent and less affected by % viral RNA, and cells with % viral RNA below that threshold were excluded from further analysis (**Supplementary Figure 4**). To enable comparisons between samples, the median number of UMIs per cell was calculated for each infection, and the UMI counts of each cell within that infection were normalized to this median value (e.g. if the total number of UMIs in a cell was 50% of that detected in the median cell, all of its UMI counts were multiplied by 2). The number of viral RNA transcripts per cell was then calculated and log_10_-transformed.

### Single-cell sorting assay for measurement of P_P_ values

Segment-specific *P_P_* values were determined as previously described for influenza A/Panama/2007/99 (H3N2) virus^19^, and as follows. 4×10^5^ MDCK cells were seeded into each well of a 6-well dish. 24 h later, cells were washed 3x with PBS and inoculated with 0.018 PFU per cell of GFHK99 WT virus and 1 PFU per cell of GFHK99 VAR_2_ virus in 250 µL of PBS. Virus was allowed to attach at 37°C for 1 h. Inoculum was then removed and cells were rinsed 3x with PBS and 2 mL of virus medium was added to the well. After 1 h at 37°C, medium was removed and cells were washed 3x with PBS and harvested by addition of Cell Dissociation Buffer (Corning). Cells were resuspended in complete medium and washed 3x with 2 mL FACS buffer (2% FBS in PBS). A final resuspension step was performed in PBS containing 1% FBS, 10 mM HEPES, and 0.1% EDTA. Cells were strained through a cell strainer cap (Falcon) and sorted on a BD Aria II cell sorter. Gating was performed to remove debris and multiplets and one event per well was sorted into each well of a 96-well plate containing MDCK monolayers at 30% confluency in 50 µl virus medium supplemented with 1 ug per mL TPCK-treated trypsin. Following the sort, an additional 50 µl of virus medium plus TPCK-treated trypsin was added to each well and plates were centrifuged at 1,800 rpm for 2 minutes to promote cell attachment. Plates were incubated at 37°C for 48 h to allow propagation of virus from the sorted cell.

RNA was extracted from infected cells in the 96-well plate using a ZR-96 Viral RNA Kit (Zymo Research) per manufacturer instructions. Extracted RNA was converted to cDNA using universal influenza primers^85^ and Maxima RT according to manufacturer instructions. After conversion, cDNA was diluted 1:4 with nuclease-free water and used as template (4 µL per reaction) for segment-specific qPCR using SsoFast EvaGreen Supermix (Bio-Rad) in 10 µl reactions, with 200 nM final primer concentration. Primers employed targeted each segment of GFHK99 WT virus, as well as the PB2 and PB1 segments of GFHK99 VAR_2_ virus. Primer sequences are listed in **Supplementary Table 3**.

Given the MOI of GFHK99 WT virus used in the experiments, an appreciable number of wells are expected to receive two or more viral genomes, and so a mathematical adjustment is needed to estimate the probability of each genome segment being delivered by a single virion. Using the relationship between MOI and the fraction of cells infected from Poisson statistics, i.e., *f* = 1 – *e*^−MOI^, the probability of the *i*th segment being present in a singly infected cell, or *P_P,i_* can be calculated from the 96-well plate using the following equation:

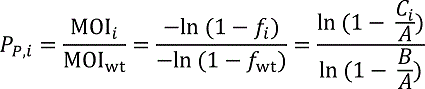

where *A* is the number of VAR_2_ wells, *B* is the number of WT wells (containing any WT segment), and *C_i_* is the number of wells positive for the WT segment in question. Wells that were negative for VAR_2_ virus segments were excluded from analysis.

