## Supplementary Information for "Collective interactions augment influenza A virus replication in a host-dependent manner"

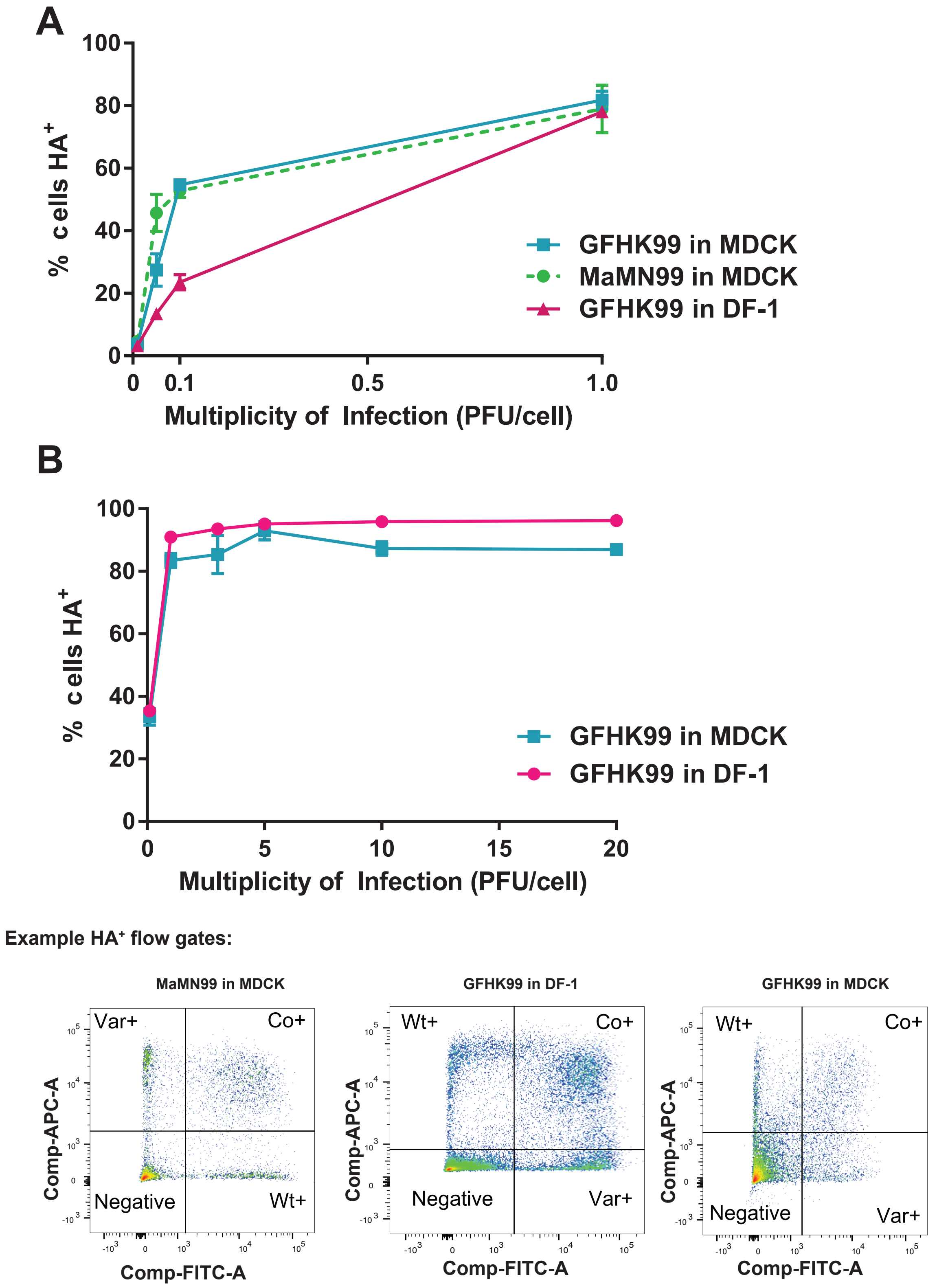


**Supplementary Figure 1 | HA positive cells detected by flow cytometry indicate levels of infection achieved across MOIs for the single cycle growth assays. (Relates to Figure 2)** Triplicate or duplicate wells of cells were harvested 24 h post infection and stained to detect surface expression of HA and HIS epitope tags. Panel A) corresponds to Figure 2 A-C and Panel B) corresponds to Figure 2 E-F. Flow gating was performed by excluding cell debris and multiplet cells. Quadrant gates were used to quantify each population.

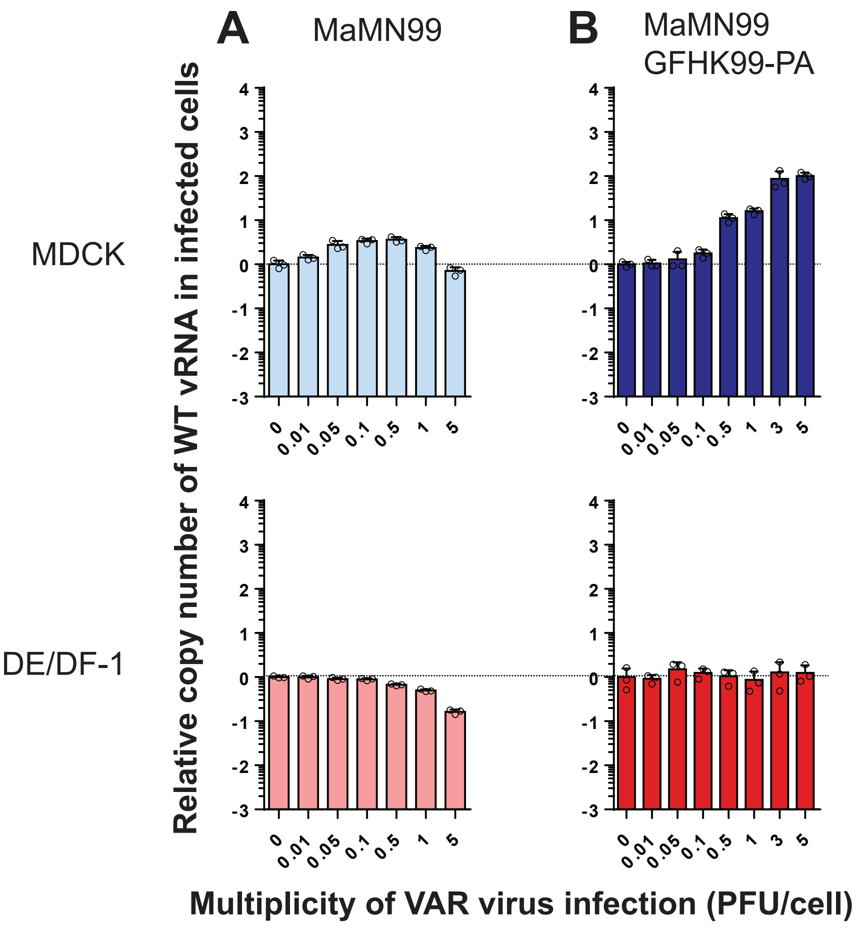


**Supplementary Figure 2 | Introduction of PA gene segment from GFHK99 virus into MaMN99 virus confers increased dependence on multiple infection for vRNA synthesis. (Relates to Figures 3 and 4)** Cells were coinfected with WT virus and increasing doses of VAR virus. WT virus MOI was 0.005 PFU per cell. The fold change in WT vRNA copy number, relative to that detected in the absence of VAR virus, is plotted for MaMN99 virus (A) and MaMN99-GFHK99-PA virus (B). Data shown in panel (A) are also shown in Figure 3. MaMN99 virus was tested in MDCK and DE cells; MaMN99 GFHK99-PA virus was tested in MDCK and DF-1 cells.

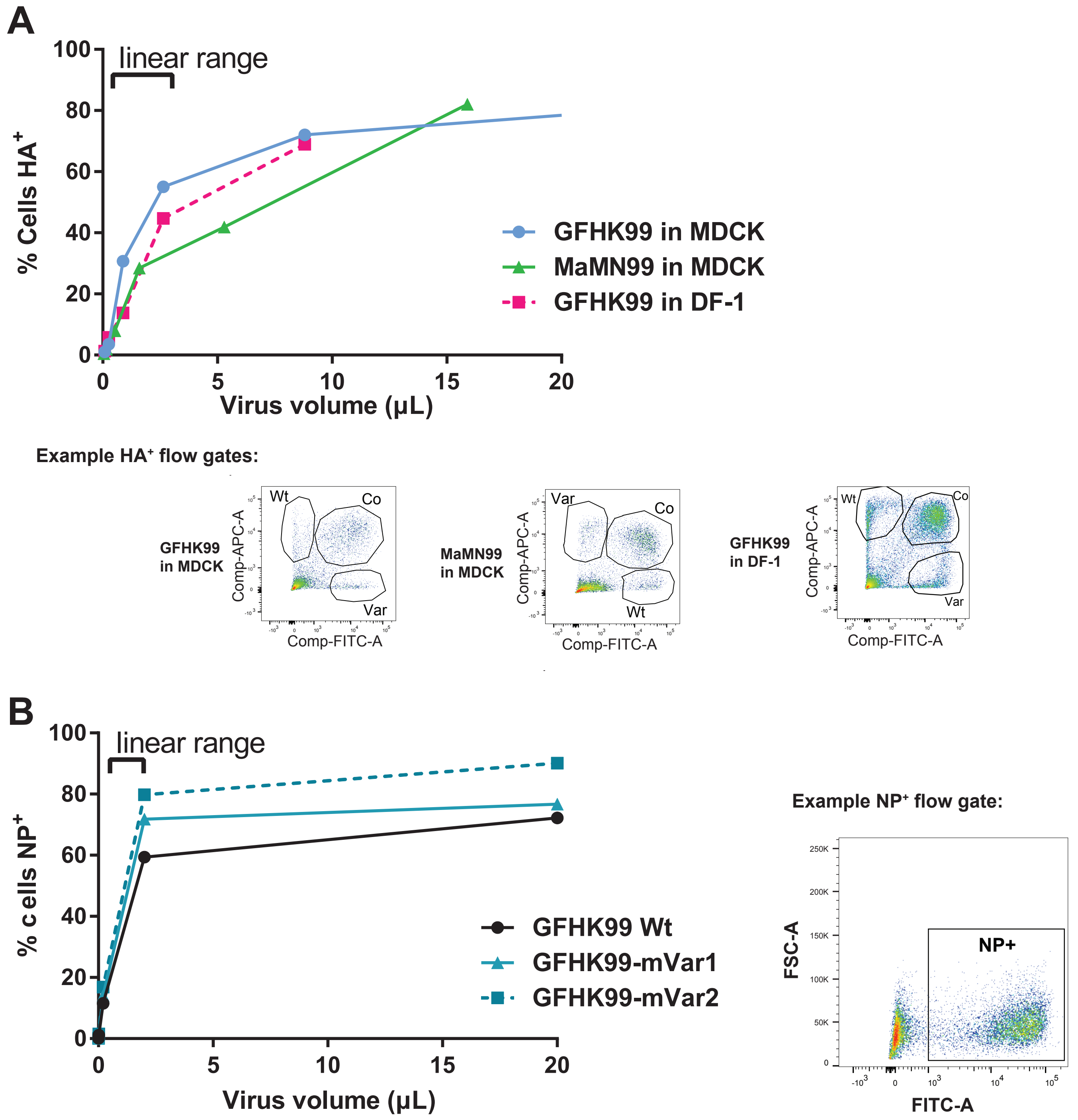


**Supplementary Figure 3 | Titration of virus stocks for HA expressing units and NP expressing units by flow cytometry. (Relates to Figures 5 and 6)** A) The doses to be used in RNA kinetics studies shown in Figure 5 were determined via flow titration of HA expressing units in the relevant cell lines. GFHK99 and MaMN99 virus mixtures were titrated in MDCK and DF-1 cell lines to calculate HA expressing units/mL in each virus-cell line combination. Serial dilutions of virus were used to infect cells under synchronized, single cycle conditions. Cells were harvested at 24 h post infection and stained for epitope tags. Data points of percent cells positive within the linear range were used to calculate the viral titer. B) GFHK99 viruses used in mRNA sequencing experiments shown in Figure 6 were titered in DF-1 cells. DF-1 cells are more permissive to infection and thus give more sensitive detection of infectious virus compared to MDCK cells. As the virus strains used did not contain epitope tags, virus detection was accomplished through cell permeabilization and detection of the viral NP protein. Data points within the linear range were used to calculate viral titers. Representative flow plots show gates used following exclusion of cell debris and doublets.

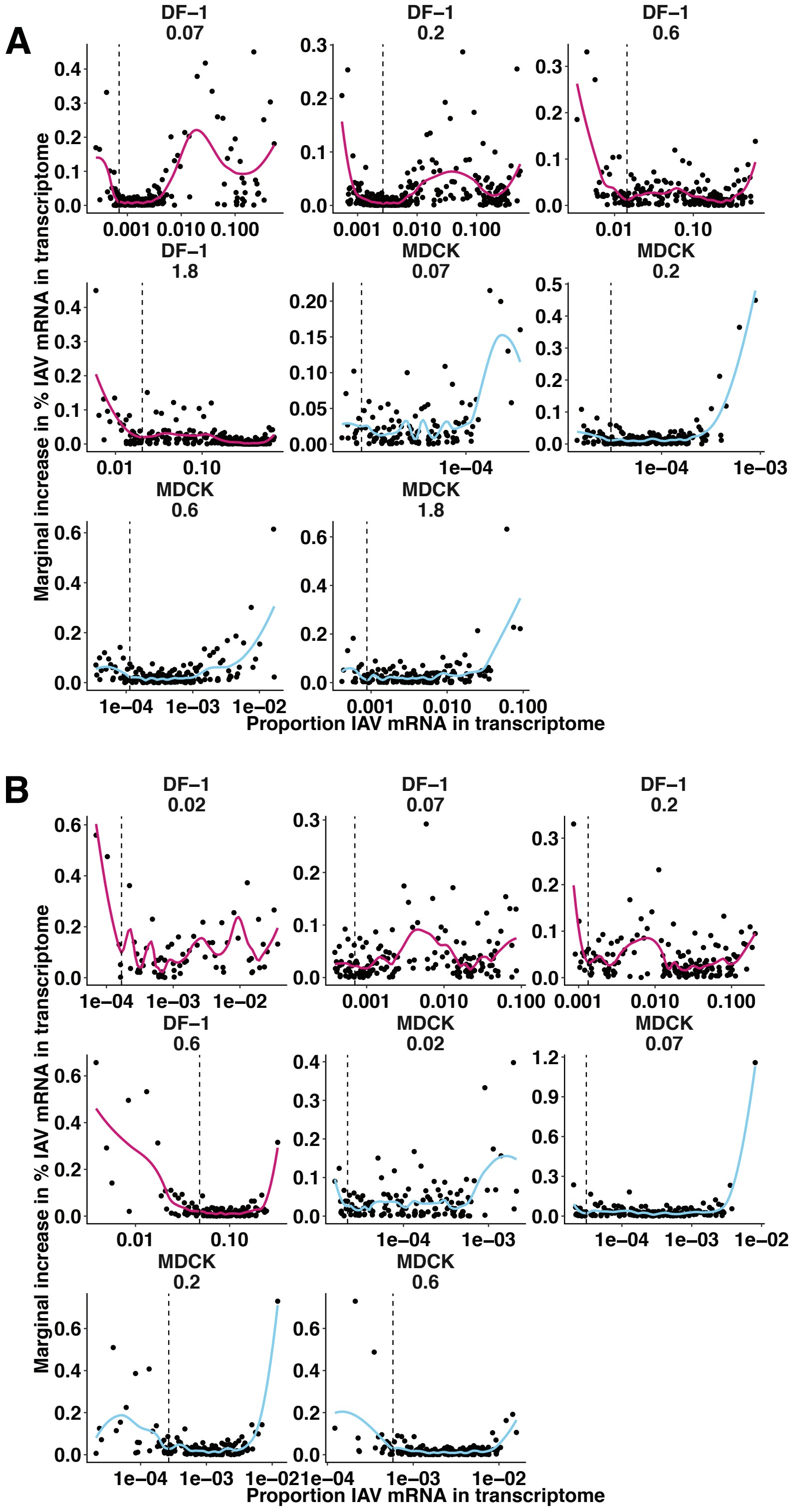


**Supplementary Figure 4 | Preliminary analysis of single-cell mRNA sequencing data to exclude cells with viral mRNA that are likely uninfected. (Relates to Figure 6)** A) Within each infection, cells in which viral RNA was detected were rank ordered by the proportion of their transcriptome that was comprised of viral RNA (% viral RNA), and the relative gain in % viral RNA from one cell to the next was plotted against the proportion of viral RNA in each cell. Local regression was performed separately for each infection, and the first local minimum of the resulting functions (indicated by dashed lines) indicated the point at which the marginal gain in % viral RNA was more consistent and less sensitive to the % viral RNA of the prior cell. Cells with % viral RNA values below this threshold were deemed falsely positive and considered uninfected for the analyses shown in **Figure 6** and **Supplementary Figure 5**. Facets indicate individual infections, with lines colored by cell type (DF-1 = pink, MDCK = blue). B) The same analysis in panel A) was applied to the data from the second experiment, in which cells were co-inoculated with a 1:1 mixture of WT and $mVAR_1$ viruses, as well as $mVAR_2$ virus at an MOI of 0.1 PFU/cell in DF-1 cells, or 1.0 PFU/cell in MDCK cells. Only cells containing all eight $mVAR_2$ segments were analyzed in this manner.

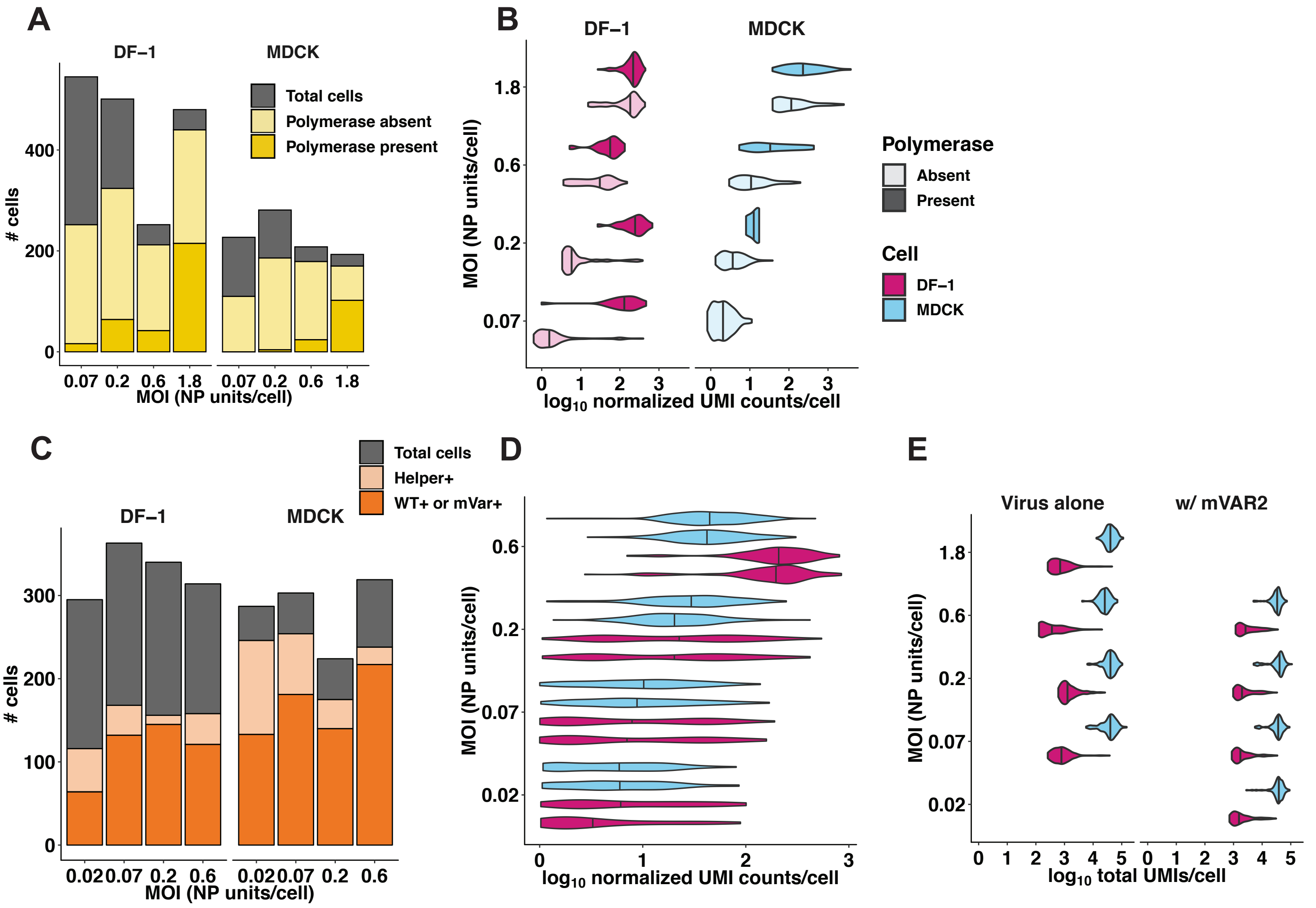


**Supplementary Figure 5 | Validation of single-cell mRNA sequencing data. (Relates to Figure 6)** A) The total number of cells sequenced, infected, and containing PB2, PB1, PA, and NP segments are represented by the cumulative heights of the gray, light yellow, and dark yellow bars, respectively. Cells that were excluded by the analysis shown in **Supplementary Figure 4** are contained within the gray bar. B) DF-1 or MDCK cells were infected with GFHK99 WT virus at four different MOIs (0.07, 0.2, 0.6, 1.8 NP units per cell), and the transcriptomes of 1,873 individual infected cells were sequenced using the 10x Genomics Chromium platform. Violin plots show distributions of $log_{10}$-transformed viral mRNA abundance, for all eight viral transcripts combined, in individual infected cells. The data are stratified by cell type (MDCK cells in blue, DF-1 cells in pink), MOI, and the presence of polymerase complex (light shading = cells missing PB2, PB1, PA, or NP; dark shading = cells in which PB2, PB1, and PA are all detected). The absence of a dark shaded distribution for MDCK cells at the lowest MOI is due to the absence of any cells in which all four of these segments were detected. C) The total number of cells sequenced, containing all eight $mVAR_2$ genome segments, and infected with either WT or $mVAR_1$ virus are represented by the cumulative heights of the gray, light orange, and dark orange bars, respectively. As in panel A), cells that were deemed falsely positive are contained within the gray bar. D) Distributions of viral UMIs per cell are shown separately for WT (bottom of each cell-MOI pair) and $mVar_1$ (top of each cell-MOI pair). Vertical lines represent the median of each distribution. E) The distributions of UMIs detected per cell are shown for each cell type, MOI, and infection type. Vertical lines represent the median of each distribution.

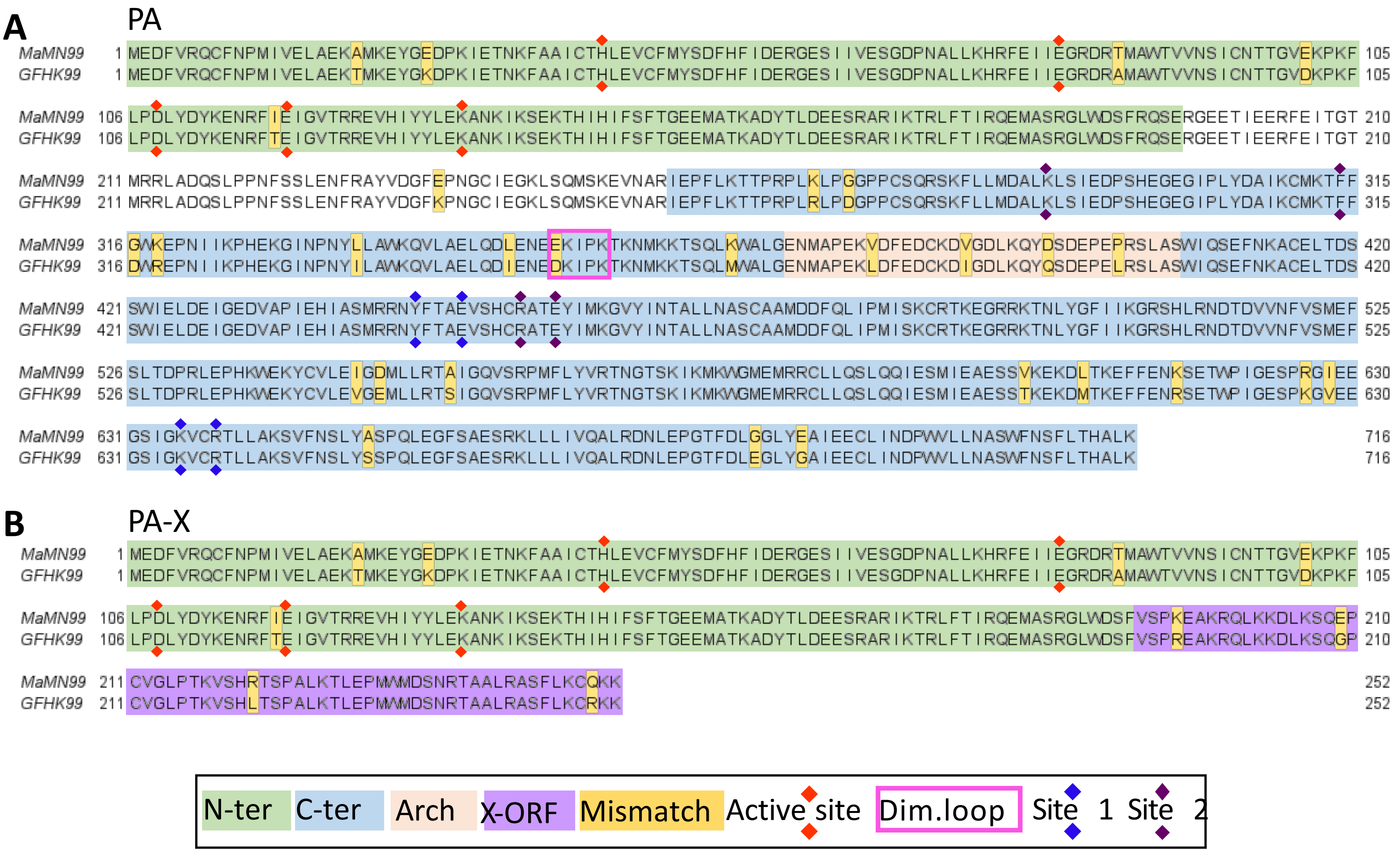


**Supplementary Figure 6 | Alignment of MaMN99 and GFHK99 virus PA and PA-X amino acid sequences.** Sequences and functional domains of the PA protein are displayed in panel (A), and those of the PA-X protein are shown in panel (B). N-ter = the N-terminal endonuclease domain[1]; C-ter = C-terminal domain[1]; X-ORF = the 61 aa region of PA-X encoded in frame 2 of the PA gene[2]; Active site = the active site of the endonuclease[3]; Dim. Loop = dimerization loop important for formation of polymerase dimers[4]; Site 1 and Site 2 = sites mediating the interaction of PA with cellular Pol II C-terminal domain[5].

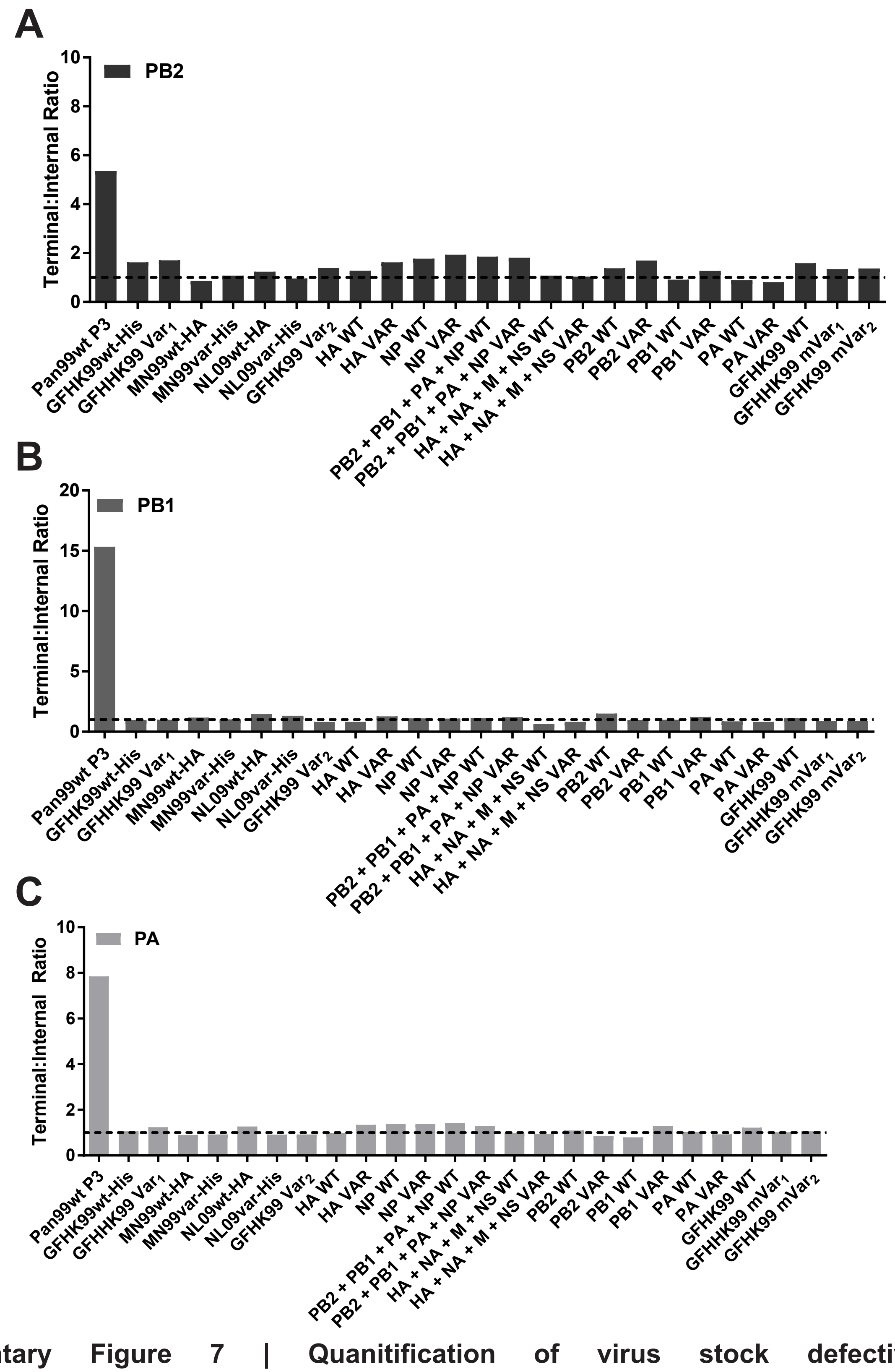


**Supplementary Figure 7 | Quanitification of virus stock defective interfering RNA content by ddPCR.** Defective RNA content for A) PB2, B) PB1, and C) PA segments was determined using primer pairs targeting terminal and internal portions of each polymerase gene segment to determine their absolute copy number and produce a ratio of terminal:internal copies. All virus stocks used in this study contained low DI content (terminal:internal ratio less than or equal to 2). A DI-rich control virus, Pan99wt P3 (A/Panama/2007/99 [H3N2]), is included for comparison. This virus stock was passaged three times in MDCK cells at high MOI. For the MaMN99-GFHK99 chimeric viruses, the segments derived from GFHK99 virus are listed in place of the full strain names.

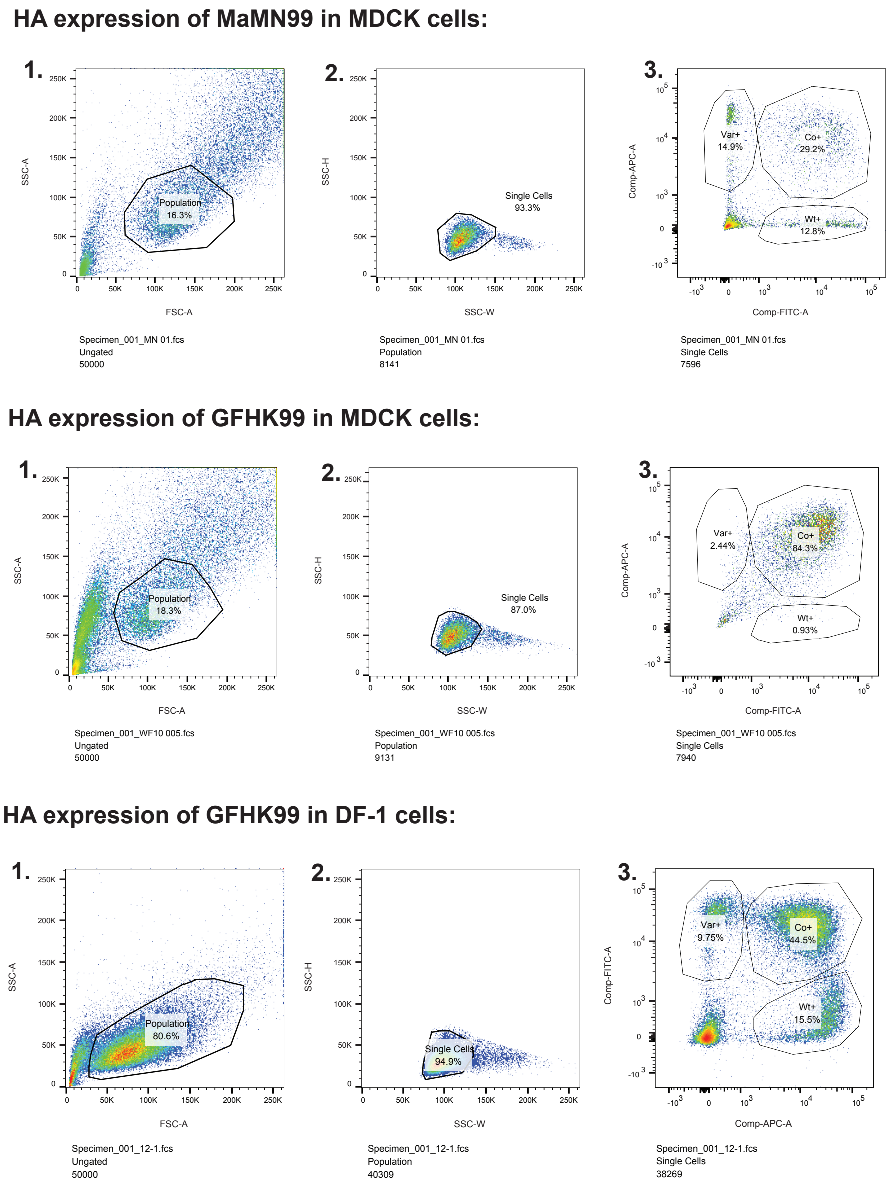


**Supplementary Figure 8 | Example gating for flow cytometry to evaluate HA positive cell numbers. (Relates to Figure 1)** Following staining for HA expression 1) a population of cells was first selected by gating out cell debris by SSC-A vs FSC-A. 2) Multiplets were excluded by gating for single cells in SSC-H vs SSC-W. In 3) populations of infected cells were gated by populations expressing the appropriate epitope tag.

### Supplementary Table 1. Genotypes of variant viruses

| | PB2 | PB1 | PA | HA | NP | NA | M | NS |
|---|---|---|---|---|---|---|---|---|
| **MaMN99[1] VAR** | G399A | G573A | G402A | A344G | A414G | G548A | A433G | A458G |
| **GFHK99[2] VAR$_1$** | A285G | A420G | A426G | T341C | T327C | T295C | A349G | T329C |
| **GFHK99-VAR$_2$** | 300G, 303T, 306C, 459C, 461A, 467T | 282C, 285C, 288G, 420G, 426C, 432T | 351G, 354T, 357T, 501G, 504T, 507T | 338G, 351C, 344C, 432G, 435A, 438T | 345G, 351A, 354G, 485C, 488A, 494A | 424G, 430A, 433A, 583G, 586C, 589C | 340A, 343G, 349G, 439A, 442T, 445G | 386T, 389A, 392G, 479G, 482C, 488G |
| **GFHK99-mVAR$_1$** | A2151G, C2164T | A2193G, A2185C | C2064A, A2061G | T1574C, G1589A | G1442A, T1411C | C1315T, G1300A | A818C, G815A | A694C, G690A |
| **GFHK99-mVAR$_2$** | A2127G, A2124G | C2175T, T2184C | C2017T, A2019G | G1553A, G1556A | G1383A, A1374G | A1255C, A1240C | C809T, T806C | A681G, C678T |
| **NL09[3] VAR** | C273T | T288C | C360T | C305T | A351G | G336A | G295A | C341T |

[1]A/mallard/Minnesota/199106/99 (H3N8), also referred to as “MN99”

[2]A/guinea fowl/Hong Kong/WF10/99 (H9N2), also referred to as “WF10”

[3]A/Netherlands/602/2009 (H1N1)

### Supplementary Table 2. Primers for the differentiation of WT and VAR by HRM

| | |
|---|---|
| **MaMN99 Primers** | |
| MN99 PB2 337 F | CCGACAACAAGCACAGTTCA |
| MN99 PB2 420 R | GCCAAAGGTCCCATGTTTTA |
| MN99 PB1 522 F | CCTCAAGGACGTGATGGAAT |
| MN99 PB1 622 R | CCATTTTCTTGGTCATGTTGTC |
| MN99 PA 379 F | GAAATTGGAGTGACACGGAGA |
| MN99 PA 461 R | TGAATGTGTGTCTTCTCGGATT |
| MN99 HA 322 F | AAACCTGGGACCTTTATGTGG |
| MN99 HA 402 R | TGAGCGATGCATAGTCTGGT |
| MN99 NP 378 F | CGACAAAGAAGAGATCAGAAGGA |
| MN99 NP 457 R | TCATCAAATGGGTGAGACCA |
| MN99 NA 522 F | TACCAGGCAAGGTTTGAAGC |
| MN99 NA 605 R | GCCCGTTACTCCAATTGTCA |
| MN99 M 404 F | TGCATGGGCCTCATATACAA |
| MN99 M 493 R | ATCAGCAATCTGCTCACACG |
| MN99 NS 389 F | GGCCATTATGGACAAGAGGA |
| MN99 NS 483 R | CGTCTGTGAAAGCCCTCAGT |
| **GFHK99[1] Primers** | |
| WF10 PB2 240 F | TGAGCAAGGCCAAACTCTTT |
| WF10 PB2 320 R | CACGTTACAGCCAGAGGTGA |
| WF10 PB1 362 F | TTGTCCAGCAAACGAGAGTG |
| WF10 PB1 441 R | AGCCGGCTGGTTTCTATTC |
| WF10 PA 386 F | GTGTGACACGGAGGGAAGTT |
| WF10 PA 461 R | TGGATATGTGTTTTCTCGGATTT |
| WF10 HA 278 F | CCCTTCTTGTGACCTGCTGT |
| WF10 HA 364 R | CCAGGGTAACACGTTCCATT |
| WF10 NP 279 F | CCTAGAGGAACATCCCAGTGC |
| WF10 NP 369 R | CAGCTCTCTCACCCATTTCC |
| WF10 NA 270 F | ATTGGTCAAAACCGCAATGT |
| WF10 NA 346 R | GCCTGCAGAAAGCCTAATTG |
| WF10 M 291 F | ACCCAAACAACATGGACAGG |
| WF10 M 373 R | TGCAACTTCCTTTGCTCCAT |
| WF10 NS 265 F | CTATCGCTTCAATGCCTGCT |
| WF10 NS 357 R | CTTTCTGCTTGGGAATGAGC |

[1]These primers were used for differentiation of GFHK99 WT and $VAR_1$ viruses.

### Supplementary Table 3. Primers and Probes for the differentiation of WT and VAR in ddPCR

| **GFHK99 WT Virus Primers** | |
|---|---|
| WF10wt PB2 286F | GACAGGGTAATGGTATCACCT |
| WF10wt PB2 480R | GGCCAGGGTTCATGTCAACCCT |
| WF10wt PB1 266F | GGTATGCACAAACAGATTGTGTAT |
| WF10wt PB1 440R | CCGGCTGGTTTCTATTCAAT |
| WF10wt PA 337F | TCTTCCGGACCTATACGACTA |
| WF10wt PA 521R | CTTCATCAAGGGTGTAGTCAG |
| WF10wt NP 336F | GAAGGAGAGACGGGAAATG |
| WF10wt NP 505R | GGCTCTTGTTCTCTGGTATG |
| WF10wt HA 323F | CGTCGAAAGATCATCAGCTGTA |
| WF10wt HA 451R | CAGGTTGTGTCTGGGAAGATT |
| WF10wt NA 413F | CTTGGGCAGGGAACCACTTTG |
| WF10wt NA 601R | CCCAGTGACACAAACATGTAAC |
| WF10wt M 328F | GAAGCTGAAGAGGGAAATGACA |
| WF10wt M 457R | AAGAGCCACTTCTGTGGTC |
| WF10wt NS 374F | CATTAGAGTGGACCAGGCA |
| WF10wt NS 499R | CCCACTATTGCTCCTTCATCT |
| **GFHK99 VAR$_2$ Virus Primers** | |
| WF10help PB2 286F | GACAGGGTAATGGTgTCtCCc |
| WF10help PB2 480R | GGCCAGGGTTCATaTCAACtCg |
| WF10help PB1 266F | GGTATGCACAAACAGAcTGcGTgT |
| WF10help PB1 440R | CCGGCTGaTTTCTgTTCAAc |
| WF10help M 331F | GCTGAAGAGAGAGATGACG |
| WF10help M 459R | CAAGAGCCACTTCCGTAGTTA |
| WF10help NS 373F | GCATTAGAGTGGATCAAGCG |
| WF10help NS 496R | ACTATTGCCCCTTCGTCC |
| **MaMN99 Virus Primers and Probes** | |
| MaMN99 NP 378 F | CGACAAAGAAGAGATCAGAAGGA |
| MaMN99 NP 457 R | TCATCAAATGGGTGAGACCA |
| MaMN99wt NP Probe | FAM-CGT(+C)[a]AA(+G)(+C)(+A)AA(+T) A(+A)TGG-IBFQ |
| MaMN99var NP Probe | HEX-CGT(+C)AA(+G)(+C)(+G)AA(+T)AATGG-IBFQ |
| **NL09 Virus Primers and Probes** | |
| NL NP 309 F | CCCTAAGAAAACAGGAGGACCC |
| NL NP 411 R | TTGGCGCCAAACTCTCCTTA |
| NLwt NP Probe | FAM-AGAC(+G)(+G)(+A)AA(+G)T(+G)GATGA-IBFQ |
| NLvar NP Probe | HEX-AGACG(+G)(+G)(+A)AGTGGATGA-IBFQ |
| **dkHK78 Virus Primers and Probes** | |
| dkHK78 NP 467 F | CCAACTTGAATGATGCCACA |
| dkHK78 NP 552 R | TCCTTGCATCAGAGAGCACA |

| | |
|---|---|
| dkHK78wt NP probe | FAM-TGC GTA CTG +G+G+A TGG AC-IBFQ |
| dkHK78var NP probe | HEX-TGC GTA +CTG +G+A+A TGG AC-IBFQ |
| **QaHK88 Virus Primers and Probes** | |
| QaHK88 NP 313 F | AAGAAAACTGGAGGCCCAAT |
| QaHK88 NP 400 R | TCCTCCTGATCTCCTCCTTG |
| QaHK88wt NP Probe | FAM-AGG A+GA +GA+T +G+GA AAA TG-IBFQ |
| QaHK88var NP Probe | FAM-AGG A+GA +GA+C +G+GA AAA TG-IBFQ |

[a] + notation indicates Locked nucleic acid (LNA) bases

### Supplementary Table 4. Primers for quantification of viral mRNA and vRNA by ddPCR

| | |
|---|---|
| **MaMN99 Reverse Transcription and PCR Primers** | |
| MaMN99 NS 552F | GGCCGTCATGGTGGCGAAT AATGCAATTGGAATCCTCAT |
| MaMN99 NS mRNAtag_dTR 13 | CCAGATCGTTCGAGTCGT TTT TTT TTT TTT TTT AGTACTAAATAAG |
| MaMN99 NS 795F | CTTGCAGGCATTGCAAC |
| MaMN99 NS 643R | CGGACTCCCCAAGCGAATCTC |
| **GFHK99 Reverse Transcription and PCR Primers** | |
| GFHK99 vRNA NS 520F | GGCCGTCATGGTGGCGAAT CCCTTCCAGGACATACTGAC |
| GFHK99 NS mRNAtag_dTR 13 | CCAGATCGTTCGAGTCGTTTTTTTTTTTTTTTTATCATTAAATAAG |
| GFHK99 NS 592R | TCATTCCATTCAAGTCCTCCGATGAG |
| GFHK99 NS 791F | CCTTTATGCAAGCCTTACAAC |
| **MaMN99 and GFHK99 Tagged PCR Primers** | |
| vRNA | GGCCGTCATGGTGGCGAAT |
| mRNA | CCAGATCGTTCGAGTCGT |
